# Anomalous networks under the multispecies coalescent: theory and prevalence

**DOI:** 10.1101/2023.08.18.553582

**Authors:** Cécile Ané, John Fogg, Elizabeth S. Allman, Hector Baños, John A. Rhodes

## Abstract

Reticulations in a phylogenetic network represent processes such as gene flow, admixture, recombination and hybrid speciation. Extending definitions from the tree setting, an *anomalous* network is one in which some unrooted tree topology displayed in the network appears in gene trees with a lower frequency than a tree not displayed in the network. We investigate anomalous networks under the Network Multispecies Coalescent Model with possible correlated inheritance at reticulations. Focusing on subsets of 4 taxa, we describe a new algorithm to calculate quartet concordance factors on networks of any level, faster than previous algorithms because of its focus on 4 taxa. We then study topological properties required for a 4-taxon network to be anomalous, uncovering the key role of 3_2_-cycles: cycles of 3 edges parent to a sister group of 2 taxa. Under the model of common inheritance, that is, when each gene tree coalesces within a species tree displayed in the network, we prove that 4-taxon networks are never anomalous. Under independent and various levels of correlated inheritance, we use simulations under realistic parameters to quantify the prevalence of anomalous 4-taxon networks, finding that truly anomalous networks are rare. At the same time, however, we find a significant fraction of networks close enough to the anomaly zone to appear anomalous, when considering the quartet concordance factors observed from a few hundred genes. These apparent anomalies may challenge network inference methods.

## 1. Introduction

Inference of the phylogenetic history of a group of species or populations is complicated by the fact that different loci may have different histories. Gene trees can conflict with each other and with the species history due to various biological processes, including incomplete lineage sorting (ILS) within each population (Maddison, 1997). When the species history is a tree, without gene flow or reticulation, ILS may cause gene trees to match the species tree topology at a lower frequency than they match some other tree topology that conflicts with the species tree. In this case, tree topologies with higher frequency than the species tree are called anomalous, and the species tree is in the “anomaly zone” (Degnan and Salter, 2005; Degnan and Rosenberg, 2006). This phenomenon has been thoroughly studied because when it occurs, a consensus or majority vote of gene trees can mislead the inference of the species tree (Kubatko and Degnan, 2007). Fortunately, when the species phylogeny is a tree, there are no anomalous rooted gene trees on 3 taxa (Pamilo and Nei, 1988) and no anomalous unrooted gene trees on 4 taxa (Allman et al., 2011). This is the basis for quartet-based species tree inference methods, which use 4-taxon subsets in gene trees, e.g. `BUCKy` (Larget et al., 2010), `ASTRAL` (Zhang et al., 2018), `SVDQuartets` (Chifman and Kubatko, 2014).

When the species phylogeny includes reticulation events such as gene flow and hybridization, it is best represented by a species network. On species networks, anomalies caused by ILS have been much less studied than on species trees. The simplest case is when all trees displayed in the species network have the same topology, such as when all reticulations involve gene flow between sister species. In that scenario, one might expect that this tree topology is represented in gene trees in equal or higher frequency than any other tree topology. However, in the presence of reticulations, this expectation is not always met, including on 3 taxa for rooted gene trees and on 4 taxa for unrooted gene trees. Several authors found examples of anomalous networks on 4 taxa displaying a unique unrooted tree topology (Solís-Lemus et al., 2016; Baños, 2019; Allman et al., 2019), whose frequency in gene trees can be lower than that of any other unrooted tree topology, even as low as 0% (Allman et al., 2023). Solís-Lemus et al. (2016) found a similar behavior on 3-taxon networks when considering rooted gene trees, and showed that the anomalously low frequency of gene trees matching the network topology may lead to misleading and inconsistent inference of the species history. This anomaly was also shown to occur under a model of continuous gene flow, with negative impact on phylogenetic inference (Long and Kubatko, 2018; Jiao et al., 2020).

Typically, networks display more than a single tree topology. One might then expect that tree topologies displayed in the network would occur with a higher frequency among gene trees than any tree *not* displayed in the species network. When this fails, we call the network “anomalous”. If a species network is anomalous, then one may expect data from gene tree frequencies to misleadingly suggest that low-frequency topologies are not displayed in the network, which may again hinder inference of the species history.

In this work, we begin by defining anomalous networks formally. We use a slightly different definition than that of Zhu et al. (2016), who focused on the large set of parental trees instead of displayed trees. Our choice captures the anomalies on 4 taxa described above, that were shown to cause misleading inference.

Second, we advance the theoretical study of anomalous networks, focusing on 4-taxon networks. These small networks are particularly relevant because many widely-used inference methods are based on 4-taxon subsets, with pseudo-likelihood or combinatorial criteria used to obtain larger networks. This approach scales better than a direct approach on the full data, and underlies `SNaQ` (Solís-Lemus and Ané, 2016), `PhyloNet-MPL` (Yu and Nakhleh, 2015), `NANUQ` (Allman et al., 2019), `PhyNEST` (Kong et al., 2022), `ADMIXTOOLS 2` (Patterson et al., 2012; Maier et al., 2022) and `TriLoNet` (Oldman et al., 2016).

On a subset of 4 taxa there are only 3 possible unrooted binary trees, whose frequencies among gene trees are called quartet concordance factors (quartet CFs). Under the coalescent model, we prove that the root of a large network is not identifiable from its quartet concordance factors, extending a result known for the class of trees and level-1 networks (Solís-Lemus and Ané, 2016; Baños, 2019). We also prove that small “sub-blobs” with a single entry and a single exit (formally defined below) are not identifiable from quartet concordance factors. This includes parallel hybrid edges that may be due to species splitting after the appearance of a geographical barrier (e.g. glacier) and merging again when the barrier disappears. These sub-blobs can also be due to sparse taxon sampling or ghost species, whose importance is now increasingly recognized (e.g. Tricou et al., 2022). Finally, we investigate the topologies of networks that can give rise to anomalies on 4 taxa. We conjecture that an anomaly requires the network to contain a 3_2_-cycle. Informally, this is an undirected cycle involving 3 nodes, one of them hybrid between the other other two, with 2 taxa descendant from the hybrid. This is the only structure previously found to cause anomalies, and is the only one causing anomalies on level-1 networks. We prove this conjecture under additional assumptions, motivated by evidence from simulations. Conversely, any network containing a 3_2_-cycle is anomalous under some set of edge parameters, for any *ρ <* 1.

We present a new recursive algorithm to calculate quartet concordance factors predicted by any phylogenetic network, regardless of its graph complexity. Importantly, this algorithm is not limited to networks of level 1, and leverages the small sample size (4 taxa) to be more efficient than general algorithms to calculate the expected frequency of a given tree topology (Yu et al., 2012). We describe our algorithm under the extended coalescent model by Fogg et al. (2023), which includes possible correlated inheritance of multiple lineages at a given locus. With no correlation, *ρ* = 0, this model is the most widely used network coalescent model, with independent lineages. With *ρ* = 1, multiple lineages of the same locus must be inherited by the same parental population, although the parent may vary across loci. Under this coalescent model of common inheritance in which each gene tree coalesces within a species tree displayed in the network (used for instance by Gerard et al., 2011; Wu, 2020; Kong et al., 2022), we prove that there are no anomalous networks: anomalies require some de-coupling of the inheritance of multiple lineages of the same locus.

Finally, using this algorithm, we quantify the frequency of anomalous 4-taxon networks using simulations under a birth-death-hybridization process and realistic parameters of speciation, extinction, and reticulation rates. We find that anomalous 4-taxon networks occur at low frequency. We also find that a more significant proportion of 4-taxon networks are sufficiently close to the anomaly zone so as to produce anomalous empirical quartet concordance factors, as observed in a sample of 300 genes, especially when the speciation rate is similar to or higher than the coalescence rate.

## 2. Notations and Definitions

### 2.1 Phylogenetic networks, displayed trees, blobs

We use standard definitions and notations for phylogenetic networks, mostly following Steel (2016) and Baños (2019). A *rooted phylogenetic network N* ^+^ on taxon set *X* is a rooted directed acyclic graph with vertices *V* = {*r*}⊔*V*_*L*_⊔*V*_*H*_ ⊔*V*_*T*_ and edges *E* = *E*_*H*_ ⊔*E*_*T*_, where *V*_*L*_ are the leaves (of out-degree 0) in bijection with *X, r* = *r*(*N* ^+^) is the root (of in-degree 0), *V*_*T*_ are the internal tree nodes of in-degree 1 and *V*_*H*_ are the hybrid nodes of in-degree greater than 1. An edge *e* = (*p, c*) is a *tree edge* (i.e. *e* ∈ *E*_*T*_) if its child *c* is a tree node or a leaf, and a *hybrid edge* otherwise (i.e. *e* ∈*E*_*H*_). Note that degree-2 nodes are allowed, and leaves are required to have in-degree exactly 1. For a node *c*, we denote by pa(*c*) the parent nodes of *c*. For a hybrid edge *e*, its *partners* are the other hybrid edges with the same child node. We use the partial order *v* ≥*w* on nodes, when *v* is an ancestor of *w*, and say that *v* is *above w* (and that *w* is *below v*). The *lowest stable ancestor* (LSA) of *Y* ⊆*X* is the lowest node that lies on all paths from the root to leaves in *Y* (Steel, 2016, p. 263). The LSA of the network, LSA(*N* ^+^), is the LSA of *X*. If *r*(*N* ^+^) = LSA(*N* ^+^), we say that *N* ^+^ is an LSA network.

The *semidirected* phylogenetic network *N* ^−^ induced by *N* ^+^ is the graph obtained by removing all edges and nodes above *s* = LSA(*N* ^+^), undirecting all tree edges and suppressing *s* if it has degree 2, as illustrated in Fig. 1 (Solís-Lemus and Ané, 2016; Baños, 2019; Xu and Ané, 2023). Following Bordewich et al. (2018), an *up-down* path in *N* ^+^ is a sequence of distinct nodes *u*_1_*u*_2_ … *u*_*n*_ with a summit node *s* = *u*_*i*_ such that *u*_*i*_ … *u*_2_*u*_1_ and *u*_*i*_ *u*_*n*−1_*u*_*n*_ are directed paths in *N* ^+^. Up-down paths in *N* ^+^ are in bijection with paths in *N* ^−^ that do not contain any *v-structure*, that is, do not have any subsequence *p*_1_*vp*_2_ such that (*p*_1_, *v*) and (*p*_2_, *v*) are both directed edges, so the up-down terminology can be used on *N* ^−^ as well (Xu and Ané, 2023). In what follows, by “network” we mean rooted or semidirected phylogenetic network, typically denoted by *N*, adding the + or − superscript only when required.

**Figure 1.**
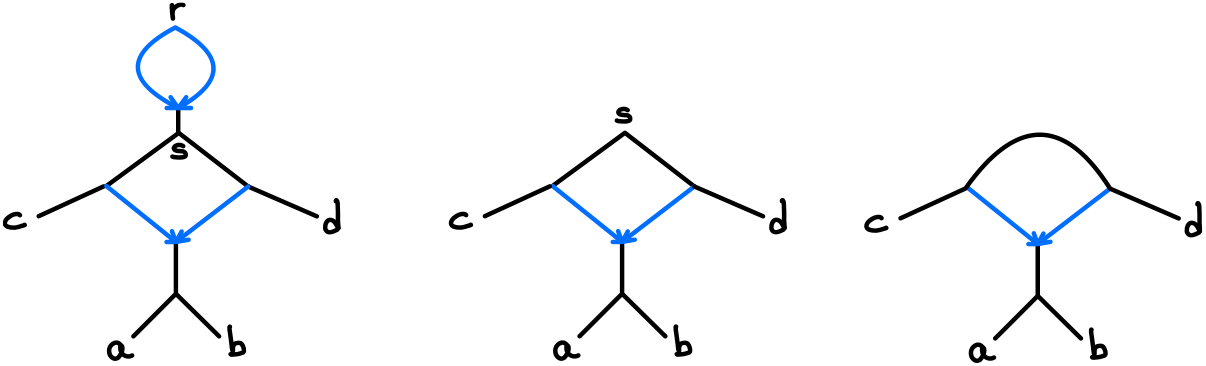
Left: rooted network *N* ^+^ whose LSA *s* is different from the root *r*, containing a 4-blob which is a 4-cycle. Middle: LSA network obtained from *N* ^+^ by removing edges and nodes above *s*, whose non-trivial blob is now a 3-blob and a 4-cycle. Right: semidirected network *N* ^−^ induced by *N* ^+^, containing a 32-cycle. Black edges are directed in *N* ^+^ and LSA(*N* ^+^), and undirected in *N* ^−^.

For a network *N* on taxa *X* and a subset *Y* ⊆ *X*, the *subnetwork N*_*Y*_ induced by *Y* is the subgraph of *N* induced by all nodes and edges on up-down paths between any two taxa in *Y* . By Baños (2019) *N*_*Y*_ is an LSA network and if *N* ^−^ is the semidirected network induced by a rooted network *N* ^+^, then 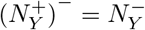

A rooted network is *binary* (or *resolved*) if all nodes in *V*_*H*_ and *V*_*T*_ have degree 3 and its root has degree A semidirected network is binary if all of its non-leaf nodes have degree 3. A network is *bicombining* if all its hybrid nodes have exactly 2 parents.

A network *N* is a *metric* network when each edge *e* is assigned a length 𝓁 (*e*) ≥ 0 and a hybridization parameter (or inheritance probability) *γ*(*e*) ∈ [0, 1] such that Σ_*u*∈pa(*v*)_ *γ*((*u, v*)) = 1 for each node *v*. In particular, *γ*(*e*) = 1 if *e* is a tree edge. Unless explicitly stated otherwise, we require that *γ*(*e*) ∈ (0, 1) if *e* is a hybrid edge.

A (topological) tree *T* is *displayed* in a network *N* if it can be obtained from *N* by deleting all but one parent hybrid edge from each hybrid node in *N*, and then deleting any edge and node not contained in any up-down path between two leaves. By default, degree-2 nodes are not suppressed, so that each edge in *T* maps to a unique edge in *N* . The set of trees displayed in *N* is denoted as 𝒟 (*N*). If *N* is semidirected, *T* is considered unrooted. When *N* is a metric network, the weight of a displayed tree *T* in *N* is defined as

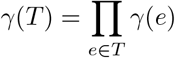

so that the weights of all trees in 𝒟 (*N*) sum to 1 (Xu and Ané, 2023).

With *U* (*T*) denoting the unrooted topology of a tree *T* after suppressing any degree-2 nodes, we say that *U* (*T*) is *induced by T* or that *T induces U* (*T*). For an unrooted tree topology 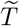 with no degree-2 nodes, its weight 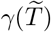 in *N* is

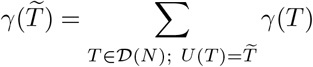

the total weight of displayed trees inducing 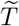 . If no tree displayed in *N* induces 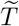, then 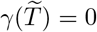.

A *blob B* in a network *N* is a maximal subgraph of *N* that is 2-edge-connected (that is, removing an edge does not disconnect it) when considered as an undirected graph. A blob is *trivial* if it consists of a single node. A node *u* in a blob *B* is a *boundary* node of *B* in *N* if there exists an edge *e* ∉ *B* incident to *B* at *u*. A blob has *degree m*, or is an *m*-blob if it is incident to *m* edges. Note that if *N* ^+^ is not an LSA network and *B* is a blob containing LSA(*N* ^+^), then the degree of *B* in *N* ^+^ is greater than the degree of the corresponding blob in the induced semidirected network *N* ^−^ (Fig. 1). Observe too that any lowest node in a non-trivial blob *B* must be a boundary and hybrid node, and every edge incident to *B* must be a cut edge. The *level* of *B* is the number of hybrid edges in *B*, minus the number of hybrid nodes in *B* (Huson et al., 2010, p. 150). If *B* is bicombining, then this is simply the number of hybrid nodes. The level of *N* is the maximum level of its blobs.

By (up-down) *cycle* in a rooted or semidirected network *N*, we mean an up-down path that forms a cycle when undirected. A *k*-cycle is a cycle with *k* edges, counted after suppressing all degree-2 nodes in *N*, except the root if *N* is rooted. A 3_2_-cycle is a 3-cycle in the semidirected network induced by *N*, containing a hybrid node *v* and two of its parents, such that *v* has at least two descendant leaves in *N* (see Fig. 3). A 3_2_-blob is a 3-blob containing a 3_2_-cycle.

**Figure 2.**
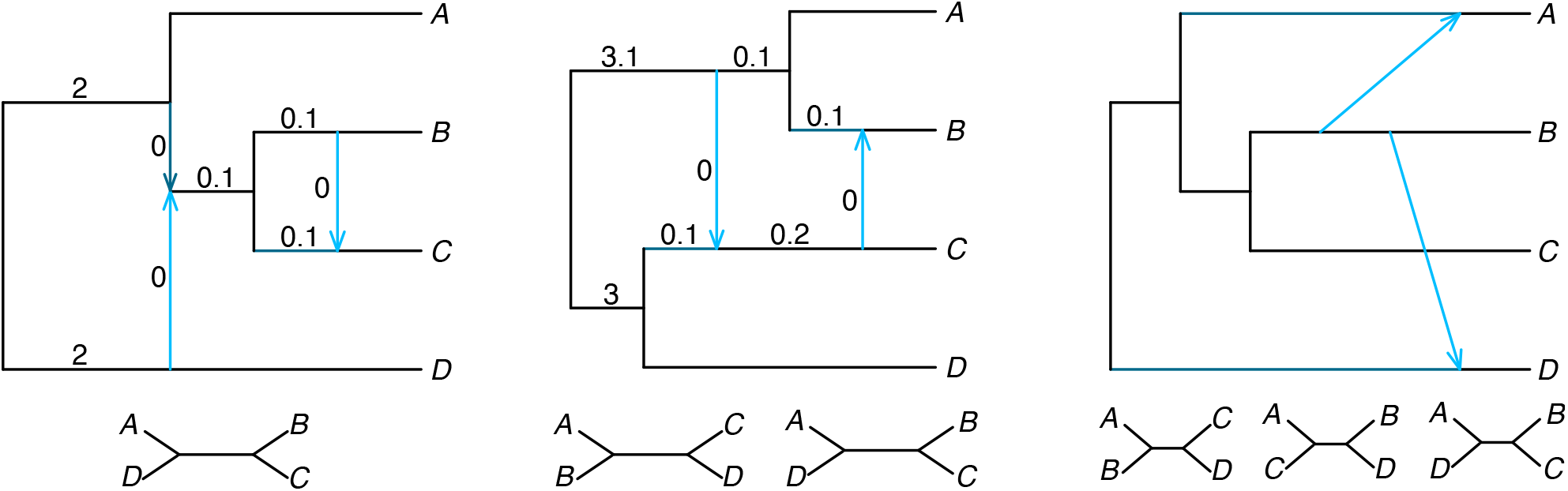
Examples of class-1 (left), class-2 (middle) and class-3 (right) networks with their displayed quartet trees (bottom). Left: With edge lengths in coalescent units as shown and all inheritance probabilities *γ* = 0.5, the network is anomalous (hard and soft) under independent inheritance, *ρ* = 0. Expected frequency of gene quartet *AD* |*BC* is 28.45%, smaller than the 35.78% of *AB* |*CD* and *AC* |*BD*. Middle: Quartet *AC* |*BD* is not displayed, but with edge lengths as shown and all *γ* = 0.5 the network is anomalous (hard and soft) if *ρ* = 0. *AC* |*BD* has frequency 24.28%, greater than the 24.11% of *AD* |*BC*. Under common inheritance, *ρ* = 1, the left and middle networks are not hard-anomalous. Right: If all *γ* = 0.5 and all edge lengths are 1, this network is soft-anomalous for all *ρ*. Its displayed quartets have weights *γ*(*AB*|*CD*) = 0.18, *γ*(*AC*|*BD*) = 0.4 and *γ*(*AD*|*BC*) = 0.42, but *AC*|*BD* is more frequent in gene trees (41.89%) than *AD*|*BC* (36.64%) for all *ρ*.

**Figure 3.**
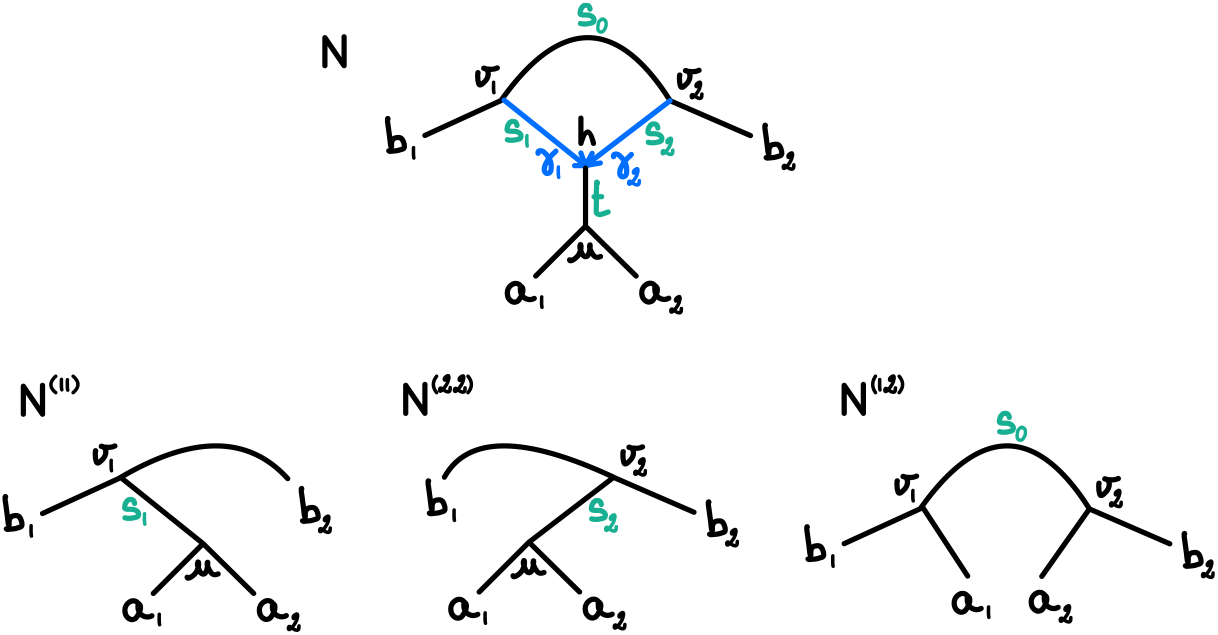
Top: semidirected network *N* with a 3_2_-cycle. Bottom: semidirected networks *N* ^(11)^, *N* ^(22)^ and *N* ^(12)^ used to calculate qCF(*N*) via (3). *N* ^(21)^ is not shown. Node labels are in black, edge lengths in green, and inheritance probabilities in blue (*γ*_1_ + *γ*_2_ = 1).

A *quartet* is an unrooted 4-taxon binary tree without any degree-2 nodes. The quartets on taxa *X* = {*a, b, c, d*} are in bijection with the non-trivial *splits* of *X*, denoted as *ab* |*cd, ac* |*bd* and *ad* |*bc*. A quartet is *displayed* in a network *N* if the corresponding 4-taxon topology is induced by a tree displayed in *N* .

### 2.2 The network coalescent and quartet concordance factors

To model the generation of gene trees on a species network, we use the coalescent model within each population, which tracks the genealogy of a sample of individuals back in time, assuming a neutral locus (Wakeley, 2008; Hahn, 2018). The coalescent time between two individuals in the same population, that is, the time when they first share a common ancestor, follows an exponential distribution with rate 1 when time is measured in coalescent units (number of generations / effective population size). With more than two individuals in a population, each pair is equally likely to be the first to coalesce.

This within-population process is used along each edge in a network. At a tree node in the species network, all gene copies are inherited from the one parent edge and the coalescent process continues along that edge, going back in time. On networks with *h >* 0 reticulations, we assume the Network Multi-Species Coalescent model with correlated inheritance NMSC(*ρ*), where *ρ* is the correlation between gene copies at reticulation events (Fogg et al., 2023). At each hybrid node and for each locus, the gene copies present at the node are inherited from one of the parental lineages according to a Dirichlet process, with the base distribution given by the inheritance probabilities *γ*(*e*), and with concentration parameter *α* = (1 −*ρ*)*/ρ*. When *ρ* = 0 for example, at each hybrid node, gene copies of the given locus have *independent* parental origins. This is the most commonly-used coalescent model assumption for species network inference (Yu et al., 2014; Solís-Lemus and Ané, 2016; Rabier et al., 2021; Allman et al., 2019; Lutteropp et al., 2022; Blair and Ané, 2020). At the other extreme, gene copies have the same parental origin when *ρ* = 1 (and this origin varies across loci). This *common* inheritance model is used by various authors, in part to improve the computational tractability of network inference (Gerard et al., 2011; Wu, 2020; Kong et al., 2022). The NMSC(*ρ*) model provides a continuum between these two extremes. It can account for different levels of gene flow across different loci, such as if selection caused some loci to be inherited from one parent more than other loci (Fogg et al., 2023).

Given a metric rooted network *N* ^+^ on taxon set *X* and a tree topology *T* on a subset *Y* ⊆*X*, rooted or unrooted, denote by ℙ_*ρ*_(*T* |*N* ^+^) the probability of *T* under the NMSC(*ρ*) model. Since these probabilities can be estimated from trees obtained from molecular alignments across multiple loci (or genes) we often refer to *T* as a “gene” tree. Similarly, we use 𝔼_*ρ*_ to denote expectation under the NMSC(*ρ*).

In particular, for a 4-taxon subset *a, b, c, d X* and an ordering of the 3 binary quartets on these taxa, we denote the expected quartet concordance factors by the vector

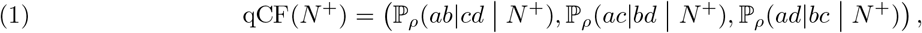

omitting the dependence on *ρ* and the four taxon names for notational convenience. Note that regardless of the value of *ρ*, the unresolved gene tree has expectation 0. In the next section (Proposition 3) we prove that qCF(*N* ^+^) is in fact fully determined by the semidirected network *N* ^−^ so that qCF(*N* ^+^) = qCF(*N* ^−^). In later sections, we consider *empirical* quartet CFs observed in a sample of gene trees: Given a sample 𝒮 = {*T*_1_,, *T*_*m*_} of *m* trees inducing binary trees on {*a, b, c, d*}, the empirical quartet CFs realized in 𝒮 the vector of quartet frequencies

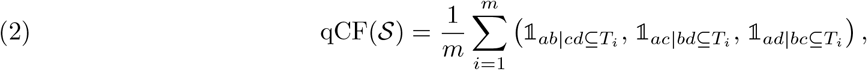

where 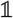 is the indicator function. In data, a gene tree may be uninformative about {*a, b, c, d*} either because it is missing one or more of these 4 taxa, or because its restriction to the 4 taxa is unresolved. Such gene trees would be ignored in calculating empirical quartet CFs, with *m* corresponding to the number of informative trees in the sample.

### 2.3 Anomalous networks

We generalize the notion of anomalous trees to networks, based on their properties under the coalescent model.

#### *Definition*1

(Anomalous network). Let *N* ^+^ be a rooted metric network on *X. N* ^+^ is *hard-anomalous* under the NMSC(*ρ*), or simply *anomalous*, if there exist unrooted binary tree topologies *T*_1_ and *T*_2_ on a taxon set *Y* ⊆ *X* such that *T*_1_ is induced by a tree displayed in 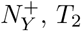, is not induced by any tree displayed in 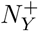, and ℙ_*ρ*_(*T*_1_|*N* ^+^) *<* ℙ_*ρ*_(*T*_2_|*N* ^+^). *N* ^+^ is *soft-anomalous* if there exist unrooted binary tree topologies *T*_1_ and *T*_2_ on *Y* ⊆ *X* such that *γ*(*T*_1_) *> γ*(*T*_2_) and ℙ_*ρ*_(*T*_1_|*N* ^+^) *<* ℙ_*ρ*_(*T*_2_|*N* ^+^).

A hard anomaly indicates a misleading signal in the sense that the highest probability gene trees are not those displayed on the network. A soft anomaly allows for a more general violation of intuition, that the hybrid parameters alone do not naively predict the ordering of gene trees by probability. Both notions capture ways in which the coalescent process on a network may lead to surprising gene tree distributions.

A hard anomaly is also a soft anomaly, because a tree topology *T* is displayed exactly when *γ*(*T*) *>* 0. This is why we refer to ‘hard-anomalous’ simply as ‘anomalous.’ Our work focuses primarily on hard anomalies because they are expected to cause more difficulties for inference. Indeed, although we chose to focus our definition on unrooted tree topologies, an analogous definition could be formulated for rooted tree topologies. Our choice is motivated by the fact that gene trees estimated from aligned molecular sequences are typically unrooted, and that error in rooted gene trees is known to have adverse effects in species phylogeny inference methods that assume gene trees are correctly rooted (Simmons and Gatesy, 2015).

Definition 1 above is similar to but different from that of Zhu et al. (2016), who focused on parental trees instead of displayed trees. Our definition extends that of Solís-Lemus et al. (2016), who first identified anomalies in networks, but restricted their study to networks that display a single quartet.

On 3 or fewer taxa, there is a single unrooted binary tree topology. On 4 taxa there are 3 unrooted binary tree topologies. We say that a 4-taxon network *N* is of *class k* if there are *k* distinct unrooted binary topologies induced by its displayed trees. An example from each class, for *k* = 1, 2, 3, is shown in Fig. 2.

While a 4-taxon network *N* of class 3 may be soft-anomalous (Fig. 2 right), it cannot be hard-anomalous because there does not exist any unrooted binary tree topologies not induced by some tree displayed in *N* . An equivalent definition of (hard-)anomalous binary networks on 4 taxa, based on their class, follows.

#### *Definition*2

(Anomalous binary networks on 4 taxa). A 4-taxon binary metric network *N* of class 1 is anomalous if ℙ_*ρ*_(*T*_1_|*N*) *<* 1*/*3 where *T*_1_ is the unique unrooted topology induced by trees displayed in *N* . A 4-taxon network *N* of class 2 is anomalous if ℙ_*ρ*_(*T*_3_|*N*) *>* min{ℙ_*ρ*_(*T*_1_|*N*), ℙ_*ρ*_(*T*_2_|*N*)} where *T*_3_ is the unique unrooted topology *not* induced by trees displayed in *N*, and *T*_1_, *T*_2_ are the other two unrooted binary tree topologies on the 4 taxa.

That Definition 2 agrees with Definition 1 follows from the next lemma.

#### Lemma 1.

*A 4-taxon binary network N displays a single quartet tree (i*.*e*., *is of class 1) if, and only if, it does not have a 4-blob*.

*Proof*. If a 4-taxon binary network *N* does not have a 4-blob, then it has two 3-blobs and either or both can be non-trivial (Allman et al., 2023, Lemma 1). Thus *N* has a cut-edge separating two pairs of taxa, whose split is the only non-trivial split in its displayed trees. (In this case, the two non-displayed quartets have equal CFs.) Conversely, let *N* be a binary network with a 4-blob. That *N* displays 2 or more quartets was proved implicitly (Allman et al., 2023, see Theorem 1 and proof) so we sketch the argument. Iteratively delete hybrid edges from *N* while it continues to have a 4-blob, until the resulting network has a lowest hybrid node *h* such that removing either parent edge of *h* leads to subnetworks without a 4-blob. Call these subnetworks *N*_1_ and *N*_2_, and let *e*_*i*_ (*i* = 1, 2) be cut-edges in *N*_*i*_ inducing non-trivial splits. Then one can show *e*_1_ and *e*_2_ correspond to incompatible splits of *N*_1_ and *N*_2_. As any tree displayed in *N*_1_ or *N*_2_ is also displayed in *N*, quartets with these splits are both displayed by *N* . □

## 3. Subgraph identifiability and subgraph requirements for anomalies

To begin our study of anomalous networks on 4 taxa, we provide some theoretical results on quartet CFs for networks on 4 or more taxa.

### 3.1 The root is not identifiable from quartet concordance factors

The following result was implicit in Xu and Ané (2023, proof of Proposition 14).

#### Lemma 2.

*Let* 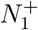 *and* 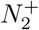 *be two metric rooted networks on X. If their induced metric semidirected networks are identical*, 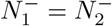, *then they have the same unrooted weighted displayed trees. In particular, the root position is not identifiable from unrooted weighted displayed trees*.

*Proof*. Let *e* be an edge in some tree displayed in 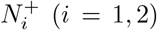. Since *e* must be on an up-down path between taxa in 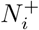, *e* cannot be above the LSA of 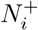 and we can assume that both 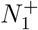 and 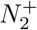 are LSA networks. Then 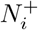 can be obtained from 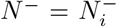 by possibly adding a degree-2 node and identifying one of the nodes as the root. Hybrid edges are then in one-to-one correspondence between 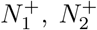 and *N* ^−^, with identical hybridization parameters. This mapping induces a weight-preserving correspondence of displayed trees on 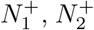 and *N* ^−^ when corresponding hybrid edges are removed from each network. The result now follows because the only difference between corresponding displayed trees is their root. □

Note that a semidirected network contains partial information about the root position in *N* ^+^. Indeed, a node *h* with incoming hybrid edges must be hybrid, and any edge incident to *h* that is not incoming must be outgoing and cannot contain the root. The direction of edges “below” *h* is also implied recursively and further constrains the possible location of the root. For example, in the semidirected network of Fig. 1 (right), the root cannot be placed on any edge incident to the parent of *a* and *b*, but could be located along any other edge.

Next we prove that, like weighted displayed trees, qCF(*N* ^+^) does not depend on the location of the root in *N* ^+^, for any value of the correlation parameter *ρ*. This is already known for level-1 networks when *ρ* = 0 (Solís-Lemus and Ané, 2016; Baños, 2019), but we prove the general case. Consequently, the root is not identifiable from quartet CFs, and whether a 4-taxon network *N* ^+^ is anomalous or not is determined by *N* ^*−*^.

#### Proposition 3.

*Let* 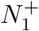 *and* 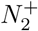 *be two metric rooted networks on a set X. If* 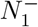 = 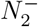, *then for every 4-taxon set* 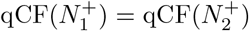. *In particular, the subgraph above the LSA of a rooted network does not affect quartet CFs*.

The proof below is inductive, to provide a roadmap for the recursive calculation of qCF presented in Algorithm 1. A shorter non-inductive proof is presented in Appendix A.

*Proof*. We need only consider the case when *X* is a set of 4 taxa. Let 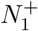 and 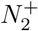 have the same semidirected network *N* ^*−*^, with *n* hybrid nodes in *N* ^*−*^. Then 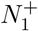 and 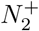 each have *n* hybrid nodes below their respective LSAs. We prove the claim by induction on *n*. Without loss of generality, we may assume that any hybrid node in 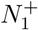 and 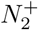 (and *N* ^*−*^) has a single child edge. If a hybrid node has two or more child edges, then we can insert a new child edge of length 0 without affecting the coalescent process, and so without changing the quartet concordance factors.

First consider *n* = 0, in which case the correlation *ρ* at hybrid nodes is not relevant. If 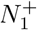 and 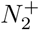 are 4-taxon trees, then the result is well known. We extend that result to allow 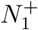 or 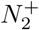 to have reticulations above their LSA, but none below the LSA. In 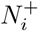, the subnetwork rooted at the LSA is a rooted tree 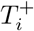, with unrooted topology *N* ^*−*^, say *a*_1_*a*_2_ *b*_1_*b*_2_ with internal edge length *t*. If *N* ^*−*^ is a star, then we resolve it to a binary tree, assigning length *t* = 0 to the new edge.

Without loss of generality, 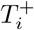 is either symmetric, or it is asymmetric with *b*_1_ sister to clade (*a*_1_, *a*_2_). Let *u*_*i*_ be the most recent common ancestor of *a*_1_ and *b*_1_ in 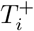. If 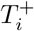 is symmetric, then *u*_*i*_ is the LSA of 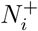 (and 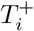). Otherwise, *u*_*i*_ is the internal node below the LSA. Let *E*_*i*_ be the event that no coalescent event occurs in 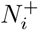 below *u*_*i*_, and 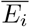 its complement. Then ℙ (*E*_*i*_) = exp(*t*) is the same for *i* = 1, 2. Conditional on *E*_*i*_, the taxa below *u*_*i*_ are exchangeable in the formation of a gene tree. These taxa are *a*_1_, *a*_2_, *b*_1_ if 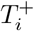 is asymmetric, all 4 taxa otherwise. Either way, the 3 quartets have equal conditional concordance factors:

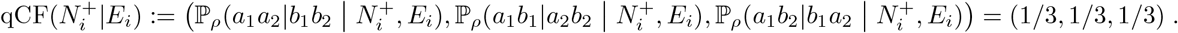

If 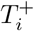 is asymmetric, then 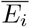 occurs when (*a*_1_, *a*_2_) coalesce along their most recent ancestral edge. If 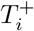 is symmetric, then 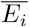 occurs when either (*a*_1_, *a*_2_) or (*b*_1_, *b*_2_) coalesce along their most recent ancestral edge. In both cases, the quartet tree matches *N* ^*−*^ conditional on 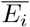,so that 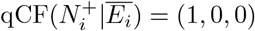 . Taken together, we find 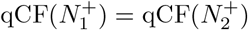.

Suppose now that there are *n ≥* 1 hybrid nodes in *N* ^*−*^. In *N* ^*−*^, choose a hybrid node *h* with no descendant hybrid node. Since hybrid nodes and edges in *N* ^*−*^ and in the LSA networks induced by 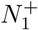 and 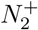 are in bijective correspondence, we reference them with identical names. The subnetwork rooted at *h* is identical in 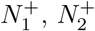 and *N* ^*−*^, and is a tree by our choice of *h*. Let *T* ^+^ denote this rooted tree and *𝓁* the number of its leaves. Note that *ρ* is not relevant to the coalescent process on *T* ^+^.

The case *𝓁* = 4 is impossible, because otherwise *h* would be above the LSA of *X*, implying *n* = 0. If *𝓁*= 3, then without loss of generality we can assume that *T* ^+^ has an edge (*h, u*) below *h*, and the subtree rooted at *u* is *a*_1_*a*_2_|*b*_1_ with internal edge length *t*. Let *E* be the event that (*a*_1_, *a*_2_) do not coalesce along this most recent ancestral edge, and 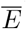 its complement. With an argument similar to the case that *n* = 0, 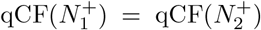 because for both networks 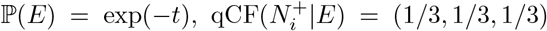 and 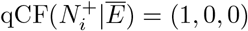 when quartet trees are ordered such that *a*_1_*a*_2_|*b*_1_*b*_2_ comes first.

For the cases *𝓁* = 1 or 2, we use induction on the network size. Let *e*_1_ = (*v*_1_, *h*), …, *e*_*m*_ = (*v*_*m*_, *h*) be the hybrid edges with child *h*, and let *γ*_*j*_ = *γ*(*e*_*j*_). Assume first that *𝓁* = 1. Let *N* ^(*j*)^ be the network obtained from 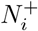 by deleting all *e*_1_, …, *e*_*m*_ except *e*_*j*_ and recursively pruning any unlabelled leaves. The tree *T* ^+^ is a single terminal edge so no coalescent event occurs on it, and

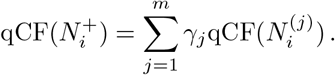

By induction, 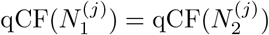 for every *j*, so 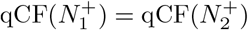.

Finally, assume 𝓁= 2. Then *T* ^+^ contains two leaves, say *a*_1_ and *a*_2_, and 3 edges: (*h, u*) of length *t*, and (*u, a*_1_), (*u, a*_2_). Let 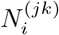 be the network obtained from 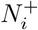 by the following steps:

1. Delete all *e*_1_, …, *e*_*m*_ except *e*_*j*_ and *e*_*k*_, and recursively prune any unlabelled leaves.
2. If *j* = *k*, contract edge (*h, u*) (or equivalently, assign it length 0).
3. If *j* ≠ *k*, delete *h, u* and their incident edges, then reconnect the network with new external (tree) edges (*v*_*j*_, *a*_1_) and (*v*_*k*_, *a*_2_), whose lengths do not affect the coalescent process.

See Fig. 3 for an example of semi-directed versions of these networks. Given a quartet ordering with *a*_1_*a*_2_|*b*_1_*b*_2_ first, we calculate qCF by conditioning on (*a*_1_, *a*_2_) coalescing along (*h, u*) or not, and if not, conditioning on the parent edge of each lineage:

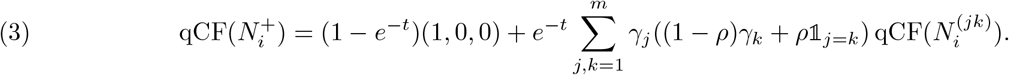

By induction, 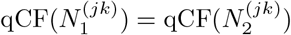 for every *j* and *k*, so 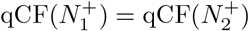 .

Note that on a network with |*X* |*>* 4, then for any subset *Y⊂ X* of 4 taxa, LSA(*Y*) can be strictly below LSA(*X*). By the second part of the proposition, the quartet CFs for *Y* are determined by the subgraph induced by all paths from LSA(*Y*) to the taxa in *Y* .

#### Example

1 (Anomaly zone for a 3_2_-cycle). We apply the formulas in the proof to a 3_2_-cycle network in Fig. 3 (top), an example of the case *𝓁*= 2 with semidirected networks *N* ^(*jk*)^ shown in Fig. 3 (bottom). Using (3) we obtain:

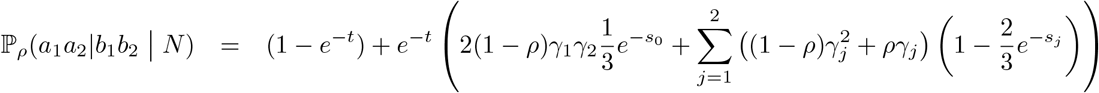

For fixed *ρ, t, s*_0_ and *γ*_1_ = 1 *− γ*_2_, this quartet CF is minimized when *s*_1_ = *s*_2_ = 0, in which case

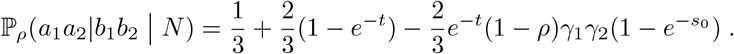

This expression generalizes eq. (1) by Allman et al. (2019) who studied the anomaly zone for a 3_2_-cycle under *ρ* = 0 (see their Fig. 5). When we set *t* = 0 and *s*_0_ tends to infinity, ℙ_*ρ*_(*a*_1_*a*_2_|*b*_1_*b*_2_ *N*) tends to 1*/*3 *−* (2*/*3)(1 *− ρ*)*γ*_1_*γ*_2_ *≤* 1*/*3. Therefore, *N* can be anomalous for any fixed value of *ρ <* 1 and any *γ*_1_ *∈* (0, 1), for sufficiently short edges incident to the hybrid node and long tree edge in the cycle.

In Example 1, the quartet CFs from a 3_2_-cycle network *N* are given by the same formulas as for a 4-taxon unrooted tree, provided the tree’s internal edge length is allowed to be negative when *N* is anomalous. Although the mechanistic coalescent process requires non-negative edge lengths, using negative branch lengths in this way will be convenient in stating subsequent results.

### 3.2 Degree-2 blobs and sub-blobs are not identifiable from quartet CFs

In this section, we show that some local subgraphs, namely *2-sub-blobs*, can be simplified into a single edge without affecting the values of quartet CFs. This is already known in a different context, for average distances on a network without the coalescent model (Xu and Ané, 2023). Here, the length of the replacement edge is defined differently, with the goal of maintaining quartet CFs specifically. We first give the 2-sub-blob definition formally.

#### Definition

3. Let *N* = (*V, E*) be a rooted or semidirected network and *G* a subgraph of *N* induced by a set *W⊂ V* of at least two vertices. Then *G* is *hybrid-closed* in *N* if, for any hybrid edge *e* in *G*, all of *e*’s partner hybrid edges in *N* are also in *G*. The *boundary ∂G* of *G* in *N* is the set of nodes *w∈ W* adjacent to a node *z ∉ W* . The graph *G* is a *sub-blob of degree 2*, or a *2-sub-blob* in *N* if it is connected, has no cut edge in *G*, is hybrid-closed in *N*, and its boundary in *N* has exactly 2 nodes. If *N* = *N* ^*−*^ is semidirected, *G traps the root* if *r*(*N* ^+^) *∈ G \∂G* for all rooted LSA networks *N* ^+^ inducing *N* ^*−*^ (e.g., Fig. 4 right).

**Figure 4.**
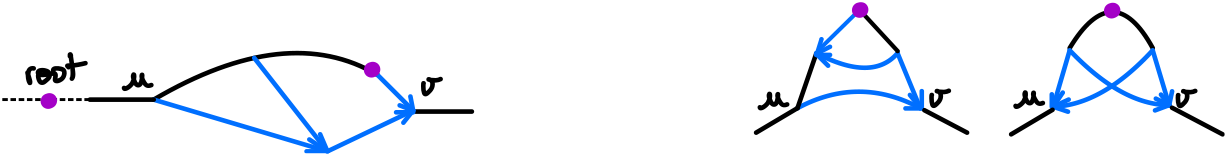
Examples of degree-2 sub-blobs *G* with boundary nodes *u* and *v* within a larger semidirected network *N* . The root may be trapped in *G* (right) or not (left).

**Figure 5.**
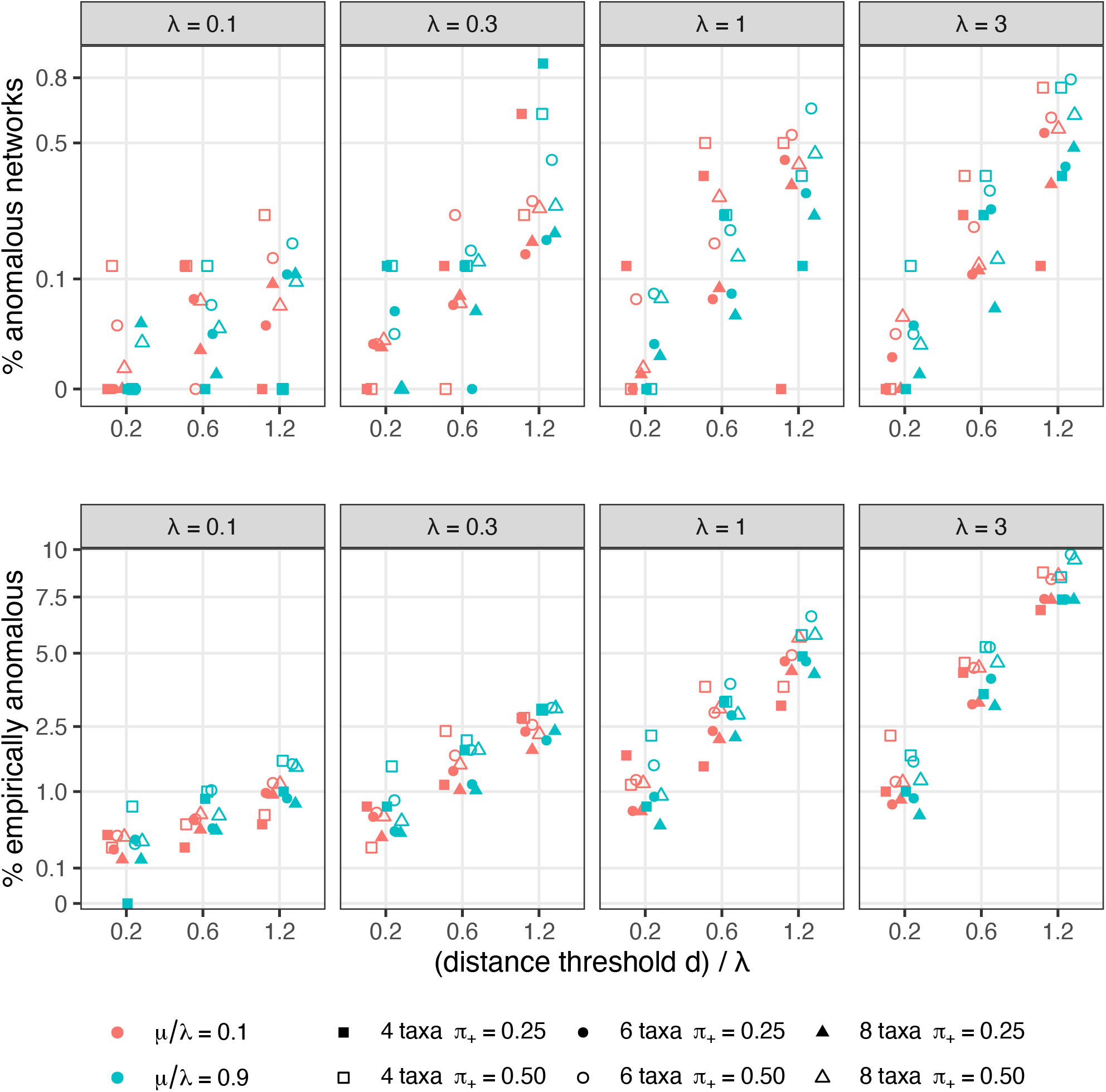
Overall percentage of anomalous 4-taxon networks among all 4-taxon subsets, under the NMSC(0). The spacing on the vertical axis corresponds to the square-root scale to better separate small values. Labels correspond to the original scale (%). Top: networks are classified as anomsalous based on their expected qCFs (1). Bottom: networks are classified based on the empirical qCFs in 300 loci (2). Simulations from different speciation rates *λ* are in different panels. Colors distinguish different extinction rates *μ* (blue: high turnover rate *μ/λ*; red: low turnover). Points are filled (resp. empty) at low (resp. high) probability of lineage-generating reticulations and high (resp. low) probability of lineage-neutral reticulations. Point shapes indicate the number of taxa in the full network: square for 4 taxa (percentage among 800 simulated networks), circle for 6 taxa (percentage among 12,000 4-taxon subnetworks), and triangle for 8 taxa (percentage among 56,000 4-taxon subnet-works).

In particular, non-trivial 2-blobs are 2-sub-blobs in a binary network *N*, as are subgraphs consisting of 2 nodes connected by parallel edges, even if they are part of a larger blob.

#### Theorem 4.

*Let N be a metric network and G be a 2-sub-blob in N with boundary nodes u and v. Then there exists t* = *t*(*G, ρ*) *≥ −* log(3*/*2) *such that replacing G with a single tree edge* (*u, v*) *or* (*v, u*) *of length t (placing the root at u or v if the root was in G) leaves the quartet concordance factors of N unchanged. If G does not trap the root, or if u (or v) has a single descendant leaf in N \ {v} (resp. N \ {u}), then t ≥* 0.

Before proving Theorem 4, we illustrate its use with an example and a corollary. Consider *G* with *m* parallel edges between two nodes *u* and *v*, of lengths *t*_*j*_ and inheritance probabilities *γ*_*j*_, *j* = 1, …, *m*. Then the *m* edges can be replaced by a single tree edge, whose length *t ≥* 0 satisfies 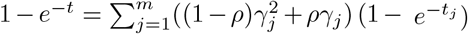 . For another application, we can rule out anomalies on simple networks:

#### Corollary 5.

*Let N be a metric 4-taxon network whose non-trivial blobs are of degree at most 2. Assume that for any 2-blob B in N with boundary nodes u and v, B does not trap the root or u (or v) has a single descendant leaf in N \ {v} (resp. N \ {u}). Then N is neither hard-nor soft-anomalous*.

*Proof*. By Theorem 4, any 2-blob in *N* can be replaced by a single edge of non-negative length, simplifying *N* to a tree. Since 4-taxon species trees (binary or star) are not anomalous, *N* is neither anomalous nor soft-anomalous.

Corollary 5 can be extended further, to networks with a 2-sub-blob *G* trapping the root, provided that it is adjacent to 2 edges (one at each boundary node) whose lengths *𝓁*_1_ and *𝓁*_2_ satisfy *𝓁*_1_ + *𝓁*_2_ *≥* log(3*/*2) *≈* 0.405 coalescent units. Then *N* has the same quartet CFs as a tree with non-negative edge lengths, and therefore is not anomalous. This follows because quartet CFs are unchanged when a degree-2 tree node *v* is suppressed, that is, when the two edges *e* and *e*^*I*^ incident to *v* (one of them a tree edge) are fused into a single edge of length *𝓁* (*e*) + *𝓁* (*e*^*I*^). Namely, let *e* = (*u, v*) be the edge replacing *G, e*_1_ = (*u, w*_1_) and *e*_2_ = (*v, w*_2_) be the edges incident to *u* and *v* in *N*, of lengths *t ≥ −* log(3*/*2), *𝓁*_1_ and *𝓁*_2_. After suppressing *v*, then *u* (and setting the root to *w*_1_ if *N* was rooted), the up-down path (*e*_1_, *e, e*_2_) is replaced by a single edge (*w*_1_, *w*_2_) of length *𝓁*_1_ + *t* + *𝓁*_2_ *≥* 0 if *𝓁*_1_ + *𝓁*_2_ *≥* log(3*/*2).

*Proof of Theorem 4*. Let *N* ^*I*^ be the network obtained from *N* after replacing *G* with a single edge of length *t* (to be determined). More specifically, all edges in *G* are removed and all nodes in *G* other than *u* and *v* are removed. A new edge *e*_new_ incident to *u* and *v* is then added. It is a tree edge in *N* ^*I*^ because in *N*, all hybrid edges going into *u* or *v* are either all in *G*, or all outside of *G*, since *G* is hybrid-closed. If *N* is rooted then we set *e*_new_ to (*u, v*) (resp. (*v, u*)) if *v* (resp. *u*) is a descendant of *u* (resp. *v*) in *N* . If neither node is a descendant of the other, then *r*(*N*) *∈ G \* { *u, v* }and we arbitrarily root *N* ^*I*^ at *u* and set *e*_new_ = (*u, v*).

We first prove that either *u* or *v* is a hybrid node of *N*, with all its parents in *G. G* must contain a hybrid node *h* because it must contain an undirected cycle. If neither *u* nor *v* is hybrid, then construct a path *p* down from *h* until *p* can no longer stay in *G*. This path *p* must end at *∂G*, say at *v*. (This is because leaves are of degree 1, so external edges are cut edges and cannot be in *G*). The last edge *e* in *p*, incident to *v*, can be removed from *G* without disconnecting *G*. Therefore, *v* is incident to another edge *e*^*I*^ in *G*. Since *p* cannot be extended in *G, e*^*I*^ must be a hybrid edge going into *v*, proving that *v* is a hybrid node with at least two parents in *G*. Since G is hybrid closed, all parents of *v* are in *G*.

Fix 4 taxa, and let *q* and *q*^*I*^ be their concordance factors in *N* and in *N* ^*I*^, for some ordering of the 3 quartets. We seek to show that *q* = *q*^*I*^ for some value of *t* still to be determined, independently of the choice of the 4 taxa. Let *H* be the random variable describing the coalescent and inheritance history of the 4 lineages along edges that are outside *G* and non-ancestral to *G*. (By coalescent and inheritance history, we mean information about edges in which coalescent events occur, which lineages coalesce at these events (Degnan and Salter, 2005) and which parental edge each lineage is inherited from at reticulations.) Since *N* and *N* ^*I*^ are identical outside *G*, the distribution of *H* is identical under *N* and *N* ^*I*^. Therefore, we need simply to show that E_*ρ*_(*q* | *H*) = E_*ρ*_(*q* ^*I*^| *H*). Let *n*_*v*_ be the number of lineages from the 4 taxa that enter *G* through *v*, when following lineages back in time. Since all edges incident to *v* that are not in *G* are children of *v*, the subnetwork rooted at *v* is outside and non-ancestral to *G*, and *H* includes all the coalescent and inheritance history in this subnetwork. Therefore *n*_*v*_ is fixed given *H*, and the distribution of *n*_*v*_ is the same under *N* and *N* ^*I*^. Since the *n*_*v*_ taxa entering *v* are exchangeable, E_*ρ*_(*q*|*H, n*_*v*_ = *j*) = E_*ρ*_(*q*^*I*^|*H, n*_*v*_ = *j*) = (1*/*3, 1*/*3, 1*/*3) for *j* = 3, 4. Thus, we may now assume *n*_*v*_*≤* 2.

First consider the case that the root of *N* can be placed at *u* or outside *G* (Fig. 4 left), and assume such a rooting for *N* . In this case, *u* is the lowest stable ancestor of *G*. Following lineages back in time, all lineages entering *G* must enter *G* through *v* and exit through *u*. If *n*_*v*_ = 0 or *n*_*v*_ = 1, then there cannot be any coalescent event between *v* and *u* (going back in time) so 𝔼_*ρ*_(*q*|*H, n*_*v*_ = *j*) = 𝔼_*ρ*_(*q*^*I*^|*H, n*_*v*_ = *j*) for *j* = 0, 1. If *n*_*v*_ = 2, we ensure that 𝔼_*ρ*_(*q*|*H, n*_*v*_ = 2) = 𝔼_*ρ*_(*q*^*I*^|*H, n*_*v*_ = 2) by setting the length *t* of *e*_new_ in *N* ^*I*^ such that

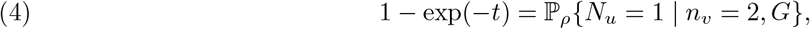

where the random variable *N*_*u*_ is the number of lineages exiting *G* at *u*. This condition ensures the probability of coalescence on (*u, v*) in *N* ^*I*^ matches that of 2 lineages entering *G* at *v* coalescing by the time they reach *u*. This probability, on the right in (4), depends on the sub-blob *G* but not on the choice of 4 taxa. It must lie in [0, 1], and therefore *t≥* 0.

Now consider the case that the root of *N* is a vertex in *G* other than *u*, which implies that *G* traps the root (Fig. 4 right). Following lineages back in time, all lineages entering *G* must enter *G* through either *v* or *u*. Since edges not in *G* are non-ancestral to all nodes of *G*, the number *n*_*u*_ of lineages entering *G* through *u* is determined by *H*. We want to show that 𝔼_*ρ*_(*q* | *H, n*_*v*_, *n*_*u*_) = 𝔼_*ρ*_(*q*^*I*^| *H, n*_*v*_, *n*_*u*_). If *n*_*u*_ + *n*_*v*_ *≤* 3, then *H* already includes a coalescence between 2 of the 4 taxa, and *q* = *q*^*I*^ is determined by *H*. If *n*_*u*_ = 3 or 4 (or similarly *n*_*v*_), then the *n*_*u*_ taxa entering *u* are exchangeable and 𝔼_*ρ*_(*q*|*H, n*_*u*_ = *j*) = 𝔼_*ρ*_(*q*^*I*^|*H, n*_*u*_ = *j*) = (1*/*3, 1*/*3, 1*/*3) for *j* = 3, 4. If *u* has a single descendant leaf in *N {v}*, then all cases are covered because *n*_*u*_ + *n*_*v*_ = 4 implies that *n*_*v*_ *≥* 3, so *G* (resp. *e*_*new*_) has no influence on *q* (resp. *q*^*I*^), and we can set *t* = 0 arbitrarily. Similarly, if *v* has a single descendant leaf in *N {u}* then all possible cases are covered and we can set *t* = 0. For the last case when *n*_*u*_ = *n*_*v*_ = 2, let *a*_1_ and *a*_2_ be the 2 taxa entering *u*, and let *b*_1_ and *b*_2_ the 2 taxa entering *v*. Since *a*_1_, *a*_2_ are exchangeable, and *b*_1_, *b*_2_ are exchangeable, 𝔼_*ρ*_(*q*|*H, n*_*u*_ = 2, *n*_*v*_ = 2) = (*x, y, y*) for an ordering of the quartets with *a*_1_*a*_2_|*b*_1_*b*_2_ first. Here *x* = 1 *−* 2*y* depends on the sub-blob *G* only, and not *H* or the choice of 4 taxa. Then 𝔼_*ρ*_(*q*|*H, n*_*u*_ = 2, *n*_*v*_ = 2) = 𝔼_*ρ*_(*q*^*I*^|*H, n*_*u*_ = 2, *n*_*v*_ = 2) provided we set *t* to be a solution of 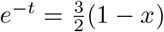 . Since *x ≥* 0, *t ≥ −* log(3*/*2) as claimed.

Although the case when the root of *N* ^+^ can be moved outside the 2-blob is easily handled in the proof above, the case when *G* traps the root is surprisingly challenging. In light of this, we state a conjecture sufficient to ensure that *t*(*G, ρ*) *≥* 0 always in Theorem 4. If this conjecture proves true, then in Corollary 5

we can remove the assumption that none of *N* ‘s non-trivial 2-blobs traps the root (as in Fig. 4 right). We obtained evidence for this conjecture through substantial simulations and numerical experiments, which did not uncover any counterexample. However, these investigations were necessarily finite and it is unclear how complicated a 2-blob might need to be to lead to an anomaly. In Appendix B, we present a proof of Conjecture 6 under additional technical assumptions.

#### Conjecture 6

*Let N* ^+^ *be a 4-taxon rooted metric network with a single non-trivial blob G, which is a 2-blob trapping the root, and with ∂G* = *u, v* . *Further suppose that a*_1_, *a*_2_ *are taxa descended from u by paths outside G, and b*_1_, *b*_2_ *taxa descended from v by paths outside G. Then* ℙ_*ρ*_(*a*_1_*a*_2_| *b*_1_*b*_2_ |*N* ^+^) *≥* 1*/*3, *so N* ^+^ *is* not *anomalous*.

### 3.3 Topological requirements for anomalies

We next seek to characterize the topology of networks that may be (hard-)anomalous for some choice of parameter values.

Binary trees are of class 1 and cannot be anomalous on 4 taxa (Larget et al., 2010; Allman et al., 2011). Level-1 networks are also well characterized: If a 4-taxon binary level-1 network *N* has a 4-cycle, then it is of class 2 and is not anomalous by applying results from Baños (2019, Proposition 6, Lemma 10). If *N* has no 4-cycle, then it is of class 1, and may have 3-cycles. With a single displayed quartet, *N* is hard-anomalous whenever it is soft-anomalous. If *N* has no 3_2_-cycle then it is not anomalous because quartet CFs on *N* are weighted averages of quartet CFs on its displayed trees. But if *N* contains a 3_2_-cycle (Fig. 3) then it is anomalous for some assignment of parameter values (Baños (2019) for *ρ* = 0; Example 1 of this article for *ρ <* 1). This anomaly occurs when the two leaves below the hybridization have a high probability of being inherited by different parents of the hybrid and when the two parents are distantly related. Then these leaves, which are sister in *N*, have a high probability of being non-sister in gene trees, each one grouping with one of the hybrid node’s sisters instead. In summary, for binary networks of level 0 or 1, anomalous networks must have a 3_2_-cycle, and we conjecture that this property holds for binary networks of any level.

#### Conjecture 7

*Let N be a 4-taxon binary semidirected metric network. If N is anomalous, then N contains a* 3_2_*-cycle*.

As evidence toward Conjecture 7, we prove it under the restriction that *N* does not have any root-trapping blob. This restriction includes the important class of networks in which there are no reticulations between the outgroup clade and the ingroup clade, and when the four-taxon set includes 1 outgroup and 3 ingroup taxa.

#### Theorem 8.

*Let N be a 4-taxon binary semidirected metric network. If N contains a* 3_2_*-cycle, then for any ρ <* 1, *N is anomalous under NMSC*(*ρ) for some set of edge lengths. As a partial converse, if N is anomalous and we further assume that N does not contain any non-trivial blob trapping the root, or if N contains a root-trapping 2-blob whose removal from N disconnects a single taxon from the other 3 taxa, then N contains a* 3_2_*-cycle as a subnetwork*.

The binary assumption in this theorem is important for ensuring that a 3-cycle is adjacent to exactly 3 edges (not part of the cycle). But it is not restrictive, because edges of length 0 are permitted. In fact, the CF of the quartet displayed in a level-1 network with a 3_2_-cycle is minimal when its hybrid edges have length 0 (Example 1). In their Proposition 1 and Fig. 6, Allman et al. (2023) construct a series of 4-taxon networks containing a growing number of nested 3_2_-cycles and show that the CF of the displayed quartet converges to the most-anomalous value of 0 (under *ρ* = 0) as the number of 3_2_-cycles increases and hybrid edge lengths are set to 0.

**Figure 6.**
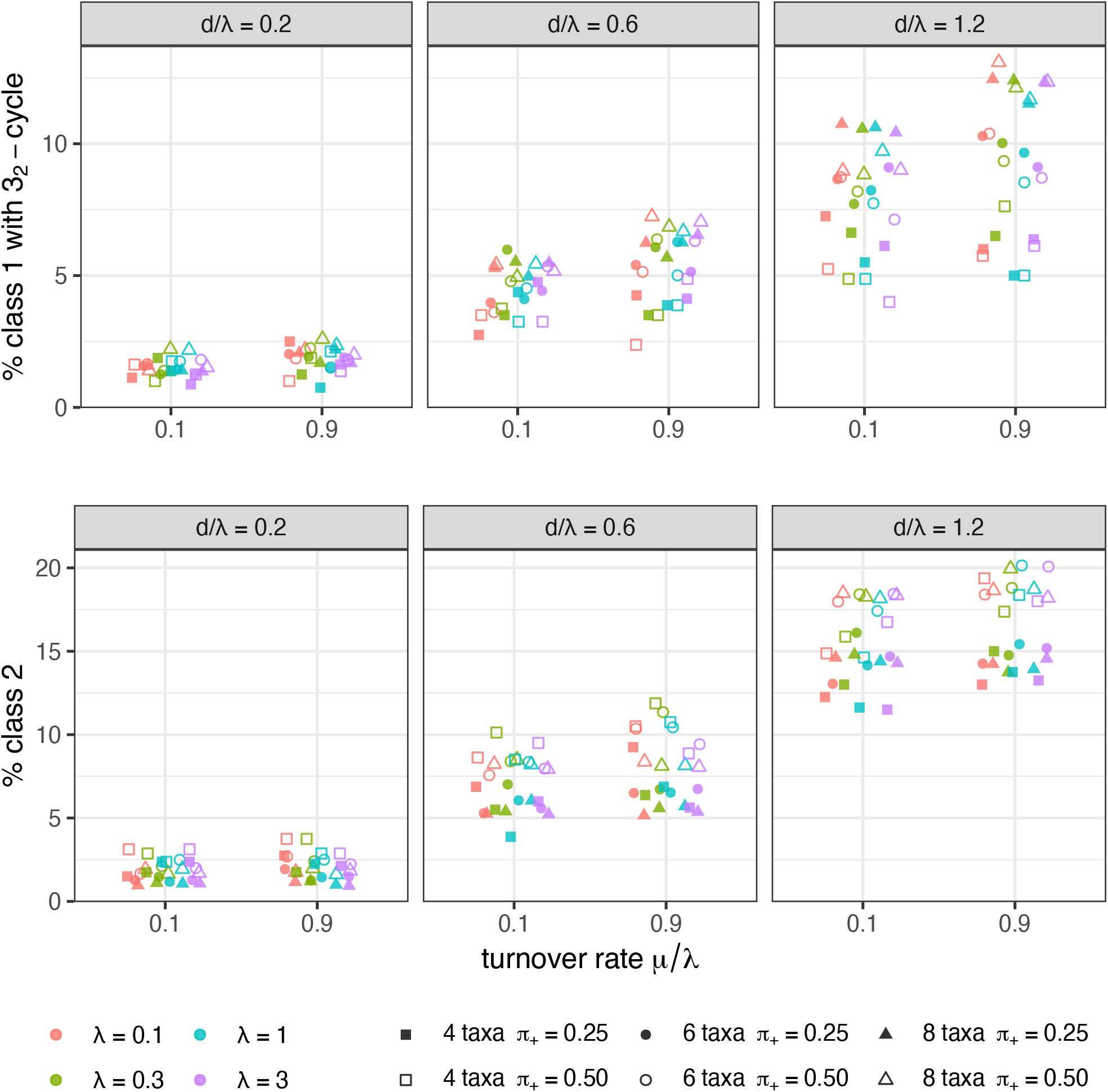
Percentage of 4-taxon networks of class 1 and containing a 3_2_-cycle (top), and of class 2 (bottom). Panels show different distance thresholds *d*, as this parameter had most influenced on the proportion of network classes. Colors distinguish the different speciation rates *λ*. The points’ fill and shape are as in Fig. 5.

The proof of Theorem 8 derives from Corollary 5 as shown below. However, if Conjecture 6 were established, it could be used in place of the weaker Corollary 5, and our argument for Theorem 8 would extend with little change to prove Conjecture 7.

We need the following useful lemmas to prove Theorem 8.

#### Lemma 9.

*For any two distinct leaves a, b of a rooted phylogenetic network N* ^+^, *there exists an up-down path from a to b in N* ^+^ *going through* LSA(*a, b*).

*Proof*. This follows from Proposition 1 of Baños (2019), which implies that the semidirected network induced by 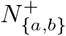 is equal to 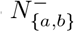 . Since *z* = LSA(*a, b*) belongs in 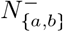, *z* is on some up-down path between two leaves in *{a, b}*, and therefore between *a* and *b*.□

#### Lemma 10.

*Any edge in a blob is in a simple up-down cycle, that is, an undirected cycle hu*_1_ … *u*_*k*_*h starting and ending at a hybrid node, such that u*_1_ … *u*_*k*_ *is an up-down path*.

*Proof*. We assume that the network is rooted, using any allowable placement of the root for a semidirected network. Let *e* = (*u, v*) be an edge in a blob. Since *e* is not a cut edge, it lies in an undirected cycle in the blob (which is not necessarily up-down). Tracing this cycle in the direction of *e*, regardless of other edge directions, there must be a last node *h* that is a descendant of *e*, since *u* is not such a descendant. The next edge in the cycle, *f*, must then be traced backward with its child node *h* being a descendant of *e* and parent node *u*^*I*^ not a descendant of *e*.

There are now down paths starting with *e* from *u* to *h*, and starting with *f* from *u*^*I*^ to *h* (having only one edge). These paths cannot edge-intersect because *u*^*I*^ is not a descendant of *e*, so *h* is hybrid. Any up paths from *u* and *u*^*I*^ to the root must intersect at or above *u* and within the blob, since they pass through its LSA. Pick any two such paths, and truncate them at their lowest intersection node, the apex. A composite path built using these paths or their reversal forms the cycle.□

#### Lemma 11.

*Let N be a 4-taxon binary phylogenetic network with a non-trivial 3-blob B. If B has a lowest hybrid node with 2 descendant taxa, then N contains a* 3_2_*-cycle*.

*Proof*. We work with a rooted network, using any permissible placement of the root if *N* is initially semidirected. Suppose *a, b* are the descendant taxa of a lowest hybrid node *h* of *B*, and *c, d* the other taxa. We may assume that *B* is the only non-trivial blob in *N* by establishing the result in the network obtained by deleting all but 1 hybrid parent edge for any hybrid node in any other non-trivial blob and then all edges with no descendant taxa. Let *M* be the subnetwork of *N* composed of all nodes and edges ancestral to *a* and *b*. Then *M* has the form of a chain of 2-blobs above the (*a, b*) cherry. Each non-trivial 2-blob in *M* is contained in *B*, but *M* may also contain trivial degree-2 nodes.

Let *z* = LSA(*c, d*). By Lemma 9, there is an up-down path *P* from *c* to *d* that passes through *z*, and thus has apex *z*. Note that *z* must be ancestral to *a, b* and in *M*, since otherwise the edge above *z* would induce the *ab* |*cd* split, and the only edges that do that are descendant edges of *h*, which are not above *c, d*. In particular, *P* passes through *M* .

Moreover, there is some up-down path from *c* to *d* that passes through a non-trivial 2-blob of *M* . To see this, consider all up-down paths from *c* to *d* that pass through *M*, and suppose none pass through a non-trivial 2-blob. Choose the lowest node *u* in *M* through which any of these paths pass. Then *u* must be a trivial 2-blob of *M*, and the edge (*u, v*) below it is a cut edge of *M* . But then in *N*, (*u, v*) is still a cut edge, which separates *a, b* from *c, d*. But by the structure of *N* there is no such node *u* above *h*, and neither *h* nor nodes below it, including *u*, are ancestral to *c* or *d*, a contradiction.

With *P* chosen to pass through a non-trivial 2-blob of *M*, consider the lowest such 2-blob it passes through. *P* contains a lowest edge which is in that 2-blob, and by Lemma 10 that edge is in an up-down cycle *C* in the blob. Let *P*_1_ be the initial segment of *P* from *c* to its first intersection with *C*, and *P*_2_ be the final segment of *P* from its last intersection with *C* to *d*. Pick a path *P*_3_ from the hybrid node of *C* down to *h. P*_3_ must not intersect with *P*_1_ or *P*_2_ since we chose the lowest 2-blob through which *P* passes and the lowest edge on *P* in that blob in picking *C*. Then *C, P*_1_, *P*_2_, *P*_3_, and the edges below *h* form a 3_2_-cycle.□

*Proof of Theorem 8*. First suppose that *N* contains a 3_2_-cycle. Then we may set *γ*(*e*) = 1 for one hybrid edge *e* in each partner pair not on the 3_2_-cycle, such that *N* is equivalent to a level-1 network with a single 3_2_-cycle as in Fig. 3. We can set edge lengths in the 3_2_-cycle known to cause an anomaly (Example 1). By continuity, *N* is also anomalous for *γ*(*e*) *∈* (0, 1) sufficiently close to 0 or 1 for hybrid edges *e* not in the 3_2_-cycle.

We turn to the second claim, and first consider networks with the property that any blob trapping the root is of degree 2 and induces a trivial split of the 4 taxa. If a network has such a blob *B*, then any other blob cannot trap the root, and *B* can be replaced by a tree edge without affecting the network’s quartet CF, by Theorem 4. Hence, we only need to consider networks whose blobs do not trap the root.

If a network is anomalous, by Corollary 5 it cannot have non-trivial blobs all of degree at most 2. Thus we only need to consider networks with at least one non-trivial 3-blob. Now consider a binary metric network *N* whose non-trivial blobs are all of degree 3 or less. Then *N* has a cut edge inducing a non-trivial split and is of class 1. Let *q*(*N*) be the CF of the quartet displayed in *N* . We show by induction on the number of hybrid nodes in *N* that *q*(*N*) *<* 1*/*3 implies the presence of a 3_2_-cycle as a subnetwork of *N* . Let *h* be a lowest (hybrid) node in a 3-blob. If *h* has 2 descendant leaves, then there exists a 3_2_-cycle in *N* by Lemma 11. If instead, *h* has a single descendant leaf, then *q*(*N*) = *γ*_1_*q*(*N*_1_) + *γ*_2_*q*(*N*_2_) where *e*_*i*_ (*i* = 1, 2) are the parent edges of *h* with inheritance probabilities *γ*_*i*_, and *N*_*i*_ is the subnetwork of *N* obtained by deleting *e*_*j*_, *j* ≠ *i* and recursively pruning any unlabelled leaves. Each *N*_*i*_ has blobs of degree 3 at most, displays the same split as *N*, has fewer hybrids than *N*, and its non-trivial blobs do not trap the root (because any cut edge that may contain the root in *N* must be retained as a cut edge in *N*_*i*_, and could contain the root of *N*_*i*_). Since *q*(*N*) *<* 1*/*3 implies *q*(*N*_*i*_) *<* 1*/*3 for at least one *i*, by induction at least one *N*_*i*_ must contain a 3_2_-cycle. This cycle is also a subnetwork of *N*, proving the claim.

Finally, consider anomalous binary networks with a 4-blob, which display at least 2 quartets by Lemma 1. As class-3 networks are never hard-anomalous, we may assume that *N* displays exactly 2 quartets, say *ab*|*cd* and *ad*|*bc*. We again use induction on the number of hybrid nodes. Let *h* be a lowest (hybrid) node of *N* in its 4-blob. It must have exactly 1 descendant, so qCF(*N*) = *γ*_1_qCF(*N*_1_) + *γ*_2_qCF(*N*_2_) with subnetworks *N*_*i*_ constructed as above. Both *N*_1_ and *N*_2_ display quartets displayed by *N*, so they cannot display *ac*|*bd*. Since qCF(*N*) is a convex sum, one of them, say *N*_1_, must be anomalous. Since the root is not trapped in a blob in *N*, the same is true for *N*_1_. Then *N*_1_ must contain a 3_2_-cycle: either by induction if *N*_1_ has a 4-blob, or by the previous steps otherwise.

#### 3_2_-*cycle containment*

While finding a network’s blobs can be done in time that is linear in the number of nodes and edges (Tarjan, 1972), it is not easy to decide if a given network with *k* hybrids contains a 3_2_-cycle. Even the number of subnetworks with exactly 1 hybrid node can be as large as *k*2^*k−*1^, growing exponentially. For a particular 4-taxon network with a 3-blob *B*, a recursive algorithm can use the following rules to decide if *B* contains a 3_2_-cycle.

1. If *B* has a lowest hybrid node with 2 descendants, then the network has a 3_2_-cycle (Lemma 11).
2. If the edges that are incident to *B* and that exit *B* (for some placement of the root) have at most 2 descendants total, then *B* does not contain a 3_2_-cycle. This is because the number of descendant leaves of a 3_2_-cycle is either 3 (when the root is outside the blob) or 4 (when the root is inside the blob).

If neither of these rules can be applied, then one can find a lowest hybrid node *h* in *B* and recursively consider the two subnetworks obtained by removing either parent edge of *h*. Although the worst-case scenario might still have an exponential time complexity, deciding if rule 1 or 2 applies takes constant time after calculating a blob decomposition, and can drastically reduce computation time in many cases.

### 3.4. Anomalies with correlated inheritance

In some networks, the dependence of quartet CFs on the correlation *ρ* can be weak. For example, quartet CFs are not affected by *ρ* at reticulations above the LSA of 3 (or all) of 4 leaves, because these leaves become exchangeable if they do not experience any coalescence before reaching their LSA. If all reticulations have a single descendant leaf, then the quartet CFs are independent of *ρ*, because there is always at most 1 individual at each hybrid node. For example, both reticulations in the class-3 network of Fig. 2 (right) have a single descendant leaf, so its quartet frequencies do not depend on *ρ*. This network is soft-anomalous for all *ρ* ∈ [0, 1]. In contrast to soft-anomalies, (hard-)anomalies cannot occur when *ρ* = 1.

#### Proposition 12.

*Under the common inheritance model with ρ* = 1, *a 4-taxon binary metric network is not anomalous*.

*Proof*. Let *N* be a 4-taxon binary network. With *ρ* = 1, each gene tree must evolve along a single tree *T* displayed in *N*, considered as a species tree, and chosen with probability *γ*(*T*). So

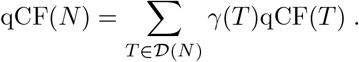

If *N* is of class 1, then all trees displayed in *N* have the same quartet topology. In each displayed tree’s qCF, the entry for the matching quartet is greater than or equal to the other two entries, so the same holds for qCF(*N*) and *N* is not anomalous.

If *N* is of class 2, let *S*_1_ and *S*_2_ denote the two quartets it displays. Order quartets with *S*_1_ and *S*_2_ first. Each tree *T* displayed in *N* displays either *S*_1_ or *S*_2_, so the third entry in qCF(*T*) is smaller than or equal to the first and second entries. Since qCF(*N*) is a convex sum, the same holds for *N*, and *N* is not anomalous.

A class-2 binary network may be soft-anomalous, not because of its non-displayed split (by Proposition 12) but because of the rankings of its displayed quartets: Rankings by *γ* weights and by CFs may differ because quartet CFs are influenced by both edge weights and edge lengths. For example, the class-2 network of Fig. 2 (middle) has two displayed splits *S*_1_ = *AB* |*CD* and *S*_2_ = *AD* |*BC* with quite different CFs under *ρ* = 1 (P_1_(*S*_1_) ≈ 0.479 and P_1_(*S*_2_) ≈ 0.316) whereas their weights are equal *γ*(*S*_1_) = *γ*(*S*_2_) = 0.5. If the inheritance probability of the upward-pointing hybrid edge *e* is increased slightly to a value above 0.5, this network becomes soft-anomalous. For example, with *γ*(*e*) = 0.51 we have *γ*(*S*_1_) = 0.49 *< γ*(*S*_2_) yet P_1_(*S*_1_) *≈* 0.474 *>* P_1_(*S*_2_) *≈* 0.319.

## 4. Recursive calculation of expected quartet concordance factors

Given a metric phylogenetic network *N*, the qCF(*N*) can be calculated by first calculating the probability of each topological gene tree, using existing software (under *ρ* = 0). However, a more efficient method of calculating them under the NMSC(*ρ*) follows the inductive proof and notation of Proposition 3. Algorithm 1 below calculates qCF(*N*) for a 4-taxon network *N* of any level, including non-binary ones.

The implementation of this algorithm in the `Julia` package `QuartetNetworkGoodnessFit` (Ané, 2023) contains several improvements to reduce complexity, which we omit from this presentation to focus on the algorithm’s main idea. First, any 2-blob with a single descendant leaf is simplified to a single edge of arbitrary length, based on Theorem 4, using that external edge lengths to not affect quartet CFs. Before this step, the root is moved to a degree-3 node, as this may move the root outside (or to the boundary) of a degree-2 blob, which may later be simplified if the 2-blob has a single descendant leaf. This is done at every iteration if recursion is needed. Second, in the case of *l* = 2 descendant leaves of a lowest hybrid node, we use the symmetry of the two “minor” CFs (line 34) to skip calculations on subnetwork *N* ^(*jk*)^ for *j > k*, so the loop on line 22 has only *m*(*m −*1)*/*2 steps instead of *m*^2^. For a bicombining network, this recursion involves 3 steps instead of 4.

In the worst-case, Algorithm 1 has exponential running time *O*(3^*h*^) where *h* is the number of hybrid nodes, assuming a bicombining network. Also, on a network with *n* taxa, this algorithm may need to be repeated for all *O*(*n*^4^) subsets of 4 taxa, so it will not scale to large networks (many taxa or many reticulations). In our experience, the shortcuts taken by cases *l* = 4 (line 11) or *l* = 3 (line 15) and by simplifying 2-blobs with a single descendant leaf lead to fast running times on many networks, including networks with number of hybrids *h ≈* 20. Further improvements could re-use calculations across different 4-taxon subsets from the same network, but the worst-case exponential time in *h* would remain.

To calculate the probability P_0_(*T* |*N* ^+^) of a rooted gene tree *T*, the generic algorithm is known to scale poorly with the degree of a blob (Elworth et al., 2019, section 13.5). In contrast, our algorithm finishes as soon as 3 taxa are below the blob regardless of the blob complexity and degree-2 blobs appearing during the recursion are eliminated as above. These improvements are specific to the context of unrooted 4-taxon gene trees.

### Algorithm 1

Recursive algorithm to calculate expected quartet CFs from a network under the NMSC(*ρ*)

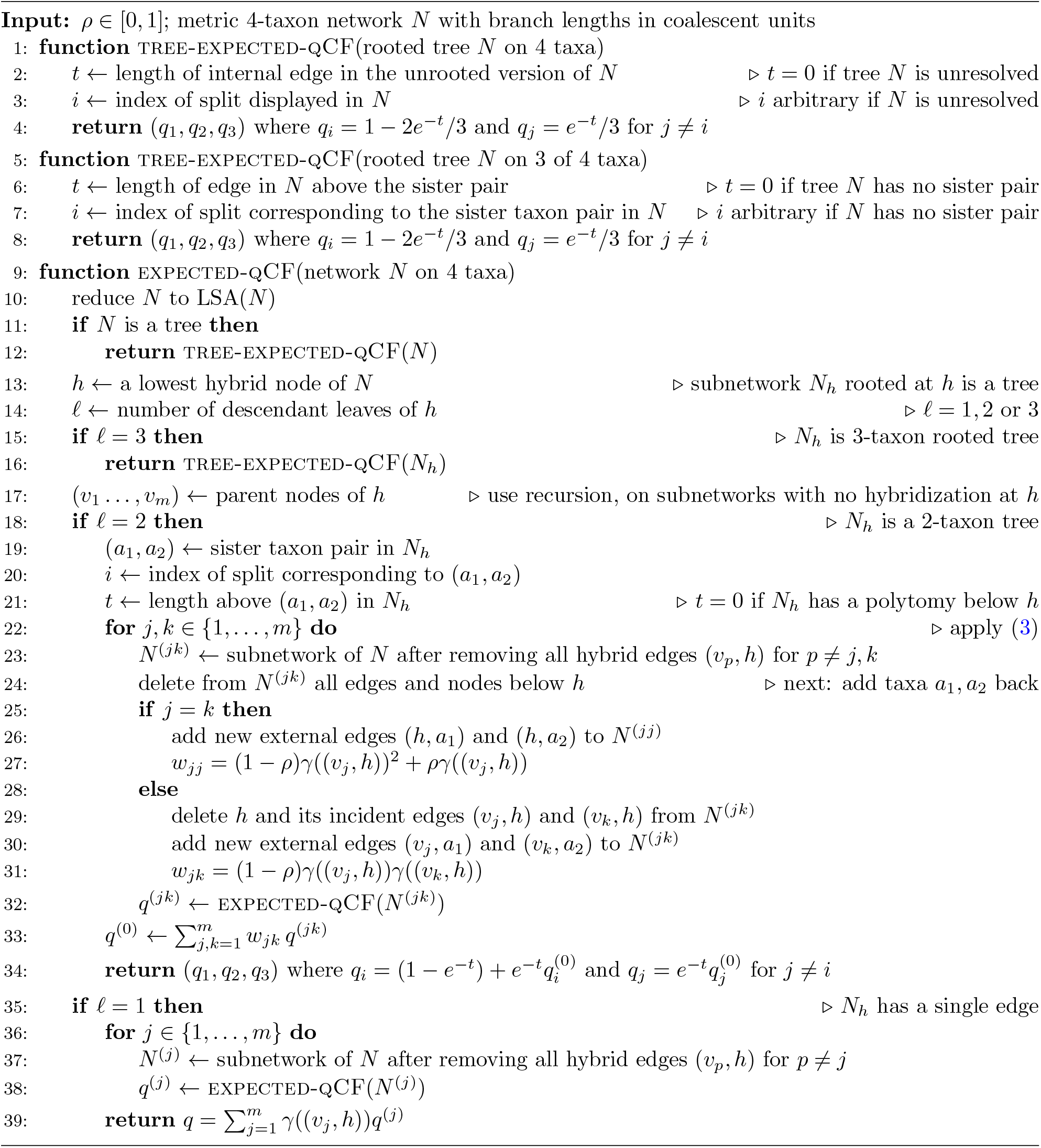

## 5. Simulation study: prevalence of hard anomalies on 4 taxa

Because anomalous networks may hinder and mislead inference from quartet CFs, we sought to quantify how often they would be encountered in practice, by simulating under a birth-death-hybridization (BDH) model with biologically realistic parameters.

### 5.1 Birth-death-hybridization networks

We simulated 800 species networks according to a BDH model first introduced by Zhang et al. (2017) as a prior for Bayesian inference of species networks. Under the BDH process, each lineage speciates with rate *λ* and goes extinct with rate *μ*. Also, each pair of coexisting lineages has rate *v* of hybridizing. We used an extension of this process as implemented in `SiPhyNetwork` (Justison et al., 2023), which allows for reticulations to depend on the genetic distance between the two involved lineages, and for three types of reticulations. Namely, when a reticulation is proposed between lineages separated by distance *δ* (calculated as the average length of all up-down paths between the two lineages in the network simulated thus far), the proposed reticulation is successful with a probability *f* (*δ*). We chose a step function with success probability 1 below a distance threshold *d*, and 0 above the threshold: *f* (*δ*) = 1 for *δ ≤d* and *f* (*δ*) = 0 if *δ > d*. Second, when a proposed reticulation is successful, it can be lineage-generating (creating a third lineage, type “+”) lineage-neutral (gene flow from one lineage into the other, type “0”) or lineage-degenerative (fusing the two lineages into a single one, type “*−*”). The probability distribution of the reticulation type, conditional on a successful reticulation, is denoted by ***π*** = (*π*_+_, *π*_0_, *π*_*−*_). Since only one event occurs at a time, the resulting networks are binary. Tree edges have positive lengths almost surely. Lineage-generating events create 2 hybrid edges of length 0, and lineage-neutral events create 1 hybrid edge of length 0 (before extinct species are pruned).

We used the general sampling strategy (GSA) as implemented in `SiPhyNetwork` to sample BDH networks with a fixed number of species *n* = 4, *n* = 6 or *n* = 8. This strategy was shown to eliminate various biases of the simple sampling strategy (Hartmann et al., 2010). Briefly, the general sampling strategy simulates networks until they have 0 or a very large number of species, then returns a network stopped at a time point when the network has the desired number *n* of species, uniformly among all times with *n* species.

Parameters for the BDH model were chosen to cover a range of values (Table 1) including speciation and hybridization rates estimated from empirical data on hominids (Bokma et al., 2012; Stadler et al., 2016), spruce, yeast (Zhang et al., 2017), and rice (Rabier et al., 2021). Reticulation-type probabilities were set to favor lineage-generating events (*π*_+_ = 0.5) or lineage-neutral events (*π*_0_ = 0.5). We chose distance thresholds *d* proportionally to the birth rate, to obtain relevant values for the total height of the simulated networks: about 14%, 44% or 90% of the median age (distance from the LSA to the tips) among simulated networks. We conducted one simulation with 144 combinations of parameters under the traditional NMSC(0) with independent inheritance (Table 1 with *ρ* fixed to 0). We conducted another simulation with 288 combinations of parameters, under the NMSC(*ρ*) with variable correlation values (Table 1 with *n* fixed to 4 taxa). For *n >* 4, each *n*-taxon network yielded 15 (for *n* = 6) or 70 (for *n* = 8) nonindependent 4-taxon subnetworks.

**Table 1.**
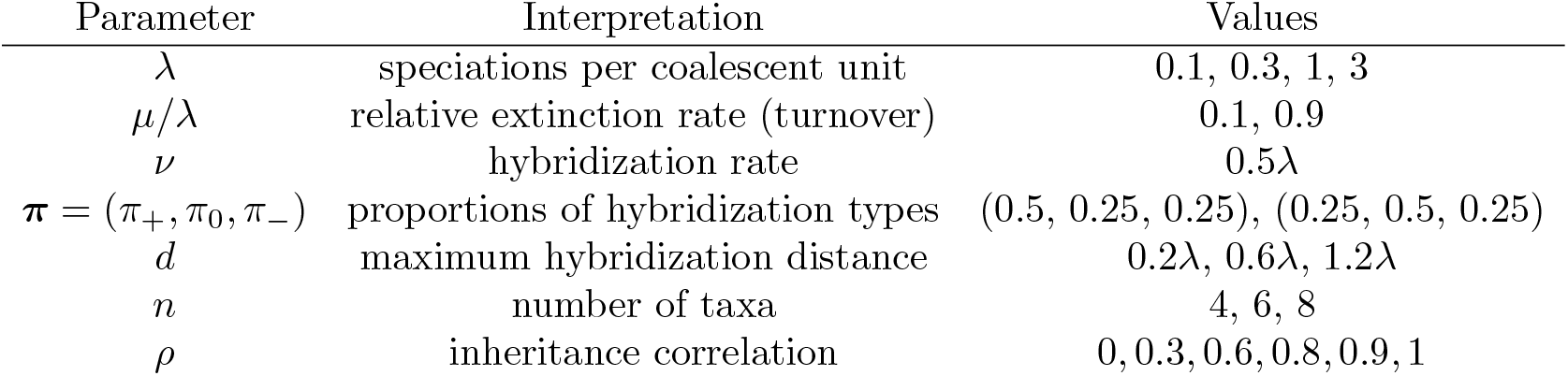
Parameters used to simulate networks under the birth-death-hybridization process. For each parameter combination, 800 networks were simulated with `SiPhyNetwork`. When *n* was varied, *ρ* was fixed to 0. When *ρ* was varied, *n* was fixed to 4. Expected quartet CFs were calculated with `QuartetNetworkGoodnessFit`. Empirical quartet CFs were simulated from 300 trees with `PhyloCoalSimulations`.

| Parameter | Interpretation | Values |
| --- | --- | --- |
| $\lambda$ | speciations per coalescent unit | 0.1, 0.3, 1, 3 |
| $\mu/\lambda$ | relative extinction rate (turnover) | 0.1, 0.9 |
| $\nu$ | hybridization rate | $0.5\lambda$ |
| $\pi = (\pi_+, \pi_0, \pi_-)$ | proportions of hybridization types | (0.5, 0.25, 0.25), (0.25, 0.5, 0.25) |
| $d$ | maximum hybridization distance | $0.2\lambda$, $0.6\lambda$, $1.2\lambda$ |
| $n$ | number of taxa | 4, 6, 8 |
| $\rho$ | inheritance correlation | 0, 0.3, 0.6, 0.8, 0.9, 1 |

`SiPhyNetwork` can simulate networks of all levels, with all types of blobs. For example, 2-blobs can result from lineage-degenerative events, or from any reticulation event followed by extinction. Also, 2-blobs naturally result from *m*-blobs with *m >* 2 when extracting the subnetwork for a subset of 4 taxa from a larger network. Justison and Heath (2022) showed that most simulated networks are complex (e.g. not tree-based) unless hybridization success requires short genetic distances between reticulating lineages.

### 5.2 Classifying 4-taxon subnetworks

From each simulated species network and each subset of 4 taxa, we extracted the subnetwork induced by the 4 taxa. To classify each 4-taxon network, we first determined its non-trivial blobs. For each network of class 1, we determined whether it had a non-trivial 3-blob. If so, we determined whether it contained a 3_2_-cycle, as we conjecture that this is a requirement for a hard-anomaly (at least under further assumptions, by Theorem 8). To do so efficiently without extracting all subnetworks, we used the algorithm described in section 3.3.

For each 4-taxon network *N* with a 4-blob, we determined its displayed quartets to determine if *N* was of class 2 or 3. In a network with *h* reticulations, there are up to 2^*h*^ displayed trees to extract to determine quartets. To avoid the computational bottleneck of this exponential complexity, we took advantage of the following. When sampling hybrid edges to extract displayed trees, we checked for the presence of 3-blobs in displayed subnetworks. If a subnetwork had a 3-blob then we did not extract its displayed trees, because all of them share the split corresponding to the subnetwork’s cut edge. This shortcut efficiently reduced the computational load.

The identification of blobs and other network manipulations to determine a network’s class were carried out with utilities from `PhyloNetworks` (Solís-Lemus et al., 2017). The `RCall` package was used to integrate `SiPhyNetwork`’s R code and Julia code (Byrne et al., 2022).

### 5.3 Detecting anomalies

For each 4-taxon subnetwork *N*, we calculated qCF(*N*) using Algorithm 1 as implemented in `QuartetNetworkGoodnessFit`. To mimic phylogenomic data sets with a few hundred unlinked loci, e.g. from target sequence capture, we simulated a sample 𝒮 of 300 gene trees according to the NMSC(*ρ*) using `PhyloCoalSimulations` (Fogg et al., 2023), and calculated the empirical qCF(𝒮) observed from these gene trees (2). Even if *N* is not anomalous, the sample of true gene trees would give an anomalous signal if qCF(𝒮) gives a greater weight to a split not displayed in the network, than to some split displayed in the network. In that case, the network would appear anomalous even in the ideal case when each gene tree can be inferred without error. We considered the network as *empirically anomalous* from 𝒮 by applying Definition 2 using the observed qCF(𝒮) instead of the theoretical qCF(𝒮).

Code to reproduce the simulation and figures is available at

https://github.com/cecileane/simulation-anomalous-4taxnet-prevalence.

### 5.4. Factors affecting the prevalence of anomalies

Overall, anomalous 4-taxon networks are rare at low speciation rates *λ* (Fig. 5). This is intuitive, because slow speciation results in long edges between speciations and, in turn, less ILS.

Anomalous networks were rare at short relative distance thresholds *d/λ* (Fig. 5). Intuitively, failure of reticulations between genetically distant populations implies fewer reticulations overall, and hence a decrease in network complexity. Indeed, the proportion of class-1 networks with a 3_2_-cycle (to exclude trees and capture class-1 networks with the potential to be anomalous) and the proportion of class-2 networks increase with the genetic distance threshold, with about 10% and 15% for each category at our highest value *d/λ* = 1.2 (Fig. 6). The prevalence of anomalous networks within each category is typically below 5-10% among class-1 networks with a 3_2_-cycle, and typically below 1-5% among class-2 networks (Fig. 7). So each category contributes about equally to the prevalence of anomalous networks overall.

**Figure 7.**
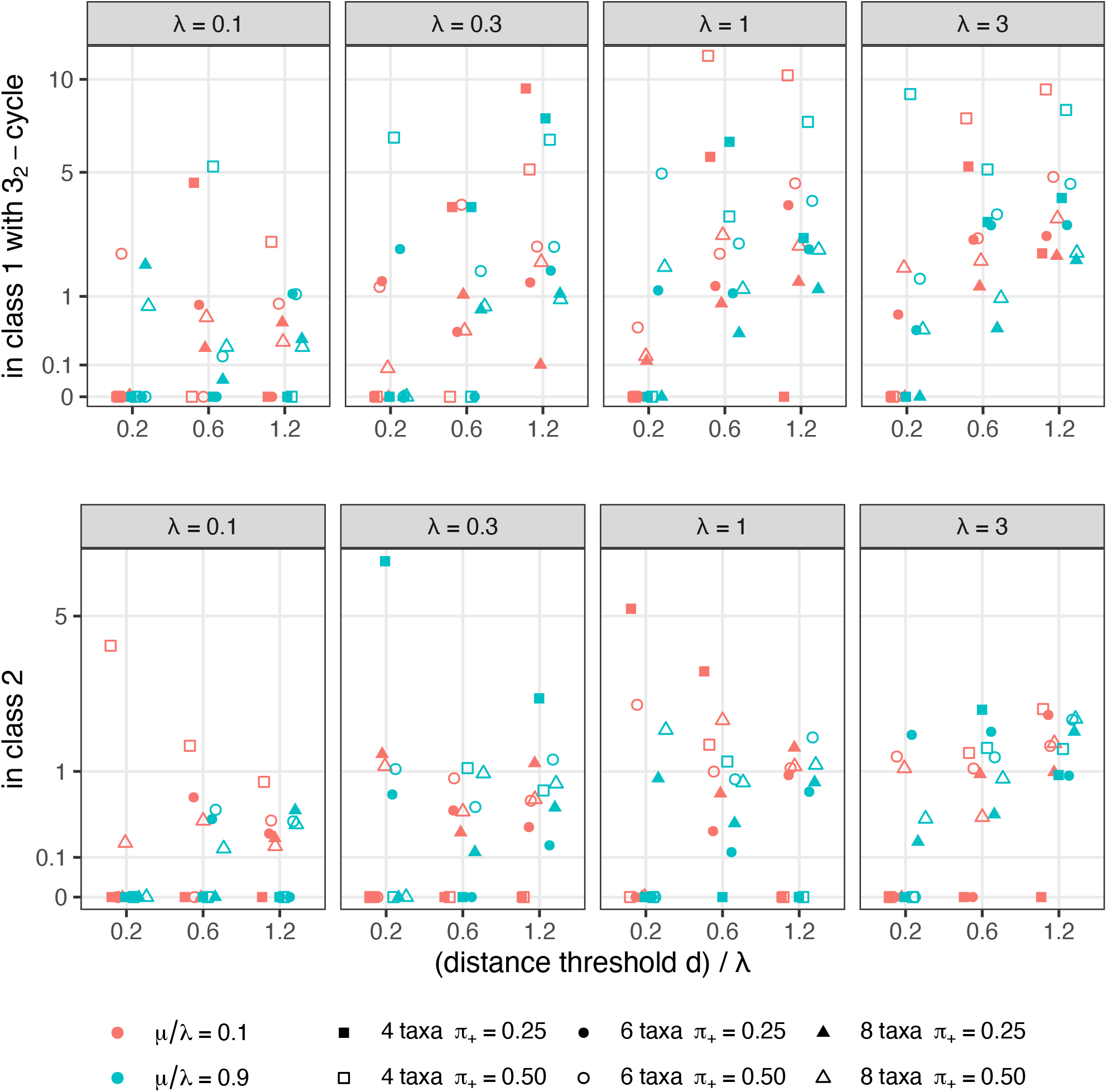
Percentage of anomalous 4-taxon networks within each subset in Fig. 6: among class-1 networks containing a 3_2_-cycle (top) and among class-2 networks (bottom). Point colors, fill and shapes are as in Fig. 5. Each point represents a variable number of 4-taxon subnetworks, most influenced by *d/λ* and the number of taxa in the full network: with an average above 200 for 6 or more taxa (circles and triangles). For 4 taxa in the full network (squares), the sample size had an average between 11 (when *d/λ* = 0.2) and 47 (when *d/λ* = 1.2) for class 1 with 3_2_-cycles, and between 20 (when *d/λ* = 0.2) and 118 (when *d/λ* = 1.2) for class 2.

The proportion of lineage-generative reticulations *π*_+_ has little influence on the proportion of the above categories (with topologies that might be anomalous), but does influence edge lengths. Lineage-degenerative and lineage-neutral events create 1 or 2 hybrid edges of positive length, while both hybrid edges have length 0 at a lineage-generative event. In a 3_2_-cycle, the most anomalous assignment of edge lengths corresponds to both hybrid edges having length 0. Therefore it is not surprising that the prevalence of anomalous networks increases with *π*_+_ (Fig. 5).

Under the wide range of parameters in our study, anomalous 4-taxon networks occur at low frequency, less than 1%. If inference methods can be robust to misleading information on 1% of 4-taxon sets, then these methods may perform well, especially at low speciation rates relative to the rate of ILS, such as appears to be the case in many biological contexts (*λ <* 0.5 in the studies cited above).

However, as many as 10% of 4-taxon networks appear empirically anomalous from a sample of 300 gene trees, at higher speciation rates like *λ* = 3 speciations per coalescent unit. This high rate means that new species form three times faster than gene lineages coalesce, and implies a high rate of ILS. Given a population size of 12,000 diploid individuals, for example, *λ* = 3 corresponds to a speciation every 8,000 generations on average. Under high diversification rates, then, the impact of anomalies could be much more pronounced and mislead inference methods, if not modeled. For example, methods assuming level-1 networks may correctly model the coalescent on an anomalous network of class 1 (with a 3_2_-cycle) but would be unable to model an anomalous network of class 2 because on 4 taxa, a level-1 4-blob cannot be anomalous.

For a non-anomalous network, an empirical anomaly may occur due to stochastic variation of the empirical qCFs around the expected qCFs. This stochastic variation would be strongest for networks with expected qCFs near the anomaly boundary, e.g. near (1*/*3, 1*/*3, 1*/*3) for class-1 networks. For a network *N* with a topology that may be anomalous (including a 3_2_-cycle), qCF(*N*) would be near the anomaly zone if its edge parameters are near parameters that make *N* anomalous. Fig. 5 suggests that a large proportion of networks are near the anomaly zone when the speciation rate and/or the distance threshold is high.

In our second simulation study with correlated inheritance, networks are never hard-anomalous when *ρ* = 1 (Proposition 12) and the prevalence of hard-anomalous networks decreases with increasing *ρ* (Figs. 8 and 9). In contrast, the prevalence of empirically anomalous networks is stable across *ρ* values. It is about 10 times higher, e.g. around 8% when *λ* = 3 and *d/λ* = 1.2 compared to 0.5-1% of true anomalies, for *ρ* = 0. When *ρ* = 1, all empirical anomalies are due to near-anomalous qCFs because none of the networks are truly anomalous. Given their large prevalence, near-anomalous networks may have a substantial practical impact on the accuracy of network inference.

**Figure 8.**
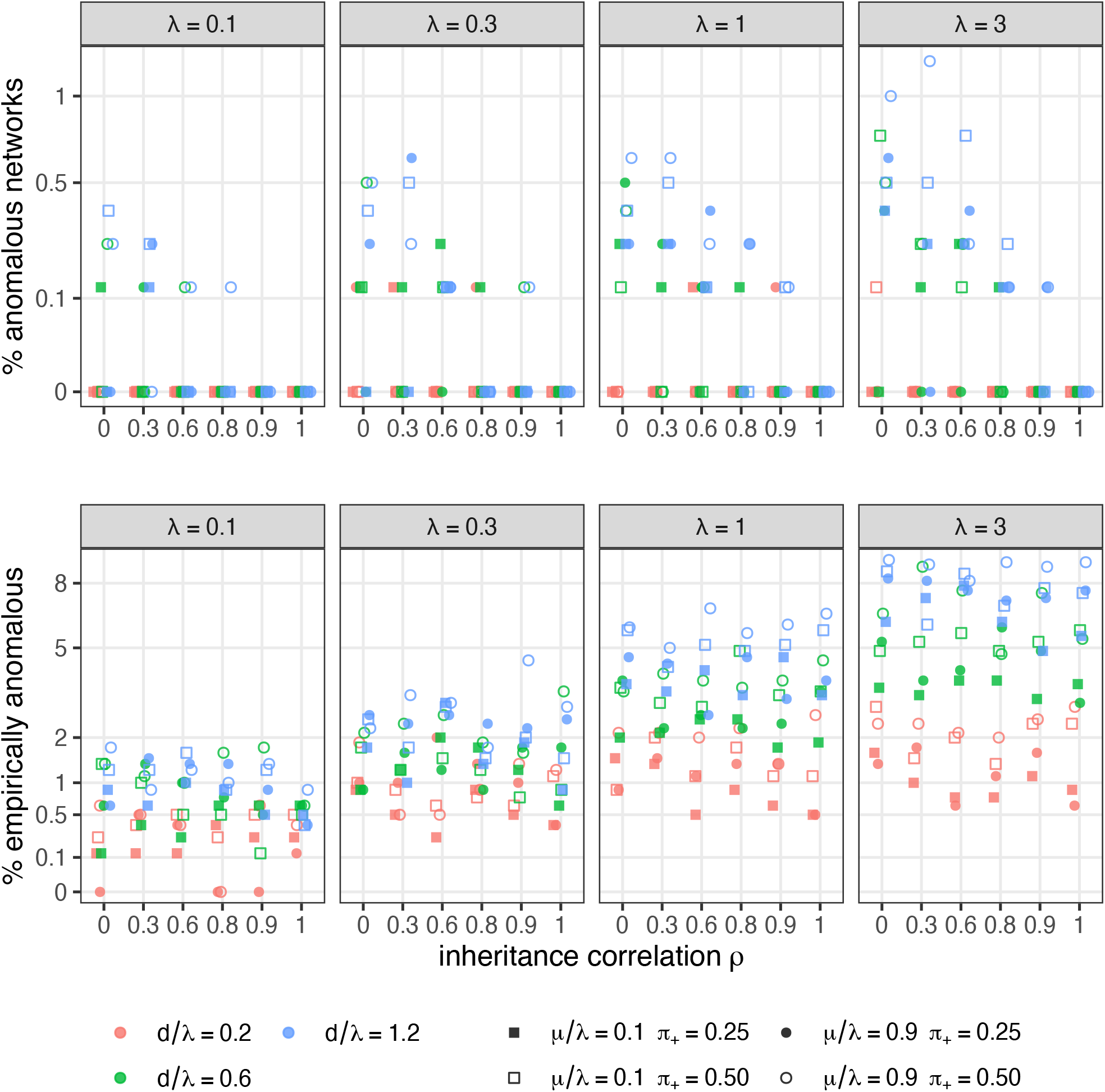
Overall percentage of anomalous 4-taxon networks among all 4-taxon subsets, under the NMSC(*ρ*) with inheritance correlation varied between 0 (independent lineages) to 1 (shared inheritance). The vertical axis is on the square-root scale. Top: networks are classified as anomalous based on their expected qCFs (Definition 2). Bottom: networks are classified based on the empirical qCFs in 300 loci. Colors distinguish the different distance thresholds *d*. Points are filled (resp. empty) at low (resp. high) probability of lineage-generating reticulations and high (resp. low) probability of lineage-neutral reticulations. Point shapes indicate the extinction rate *μ*: square for low turnover (*μ/λ*) and circle for high turnover. Each point represents the percentage among 800 four-taxon networks.

**Figure 9.**
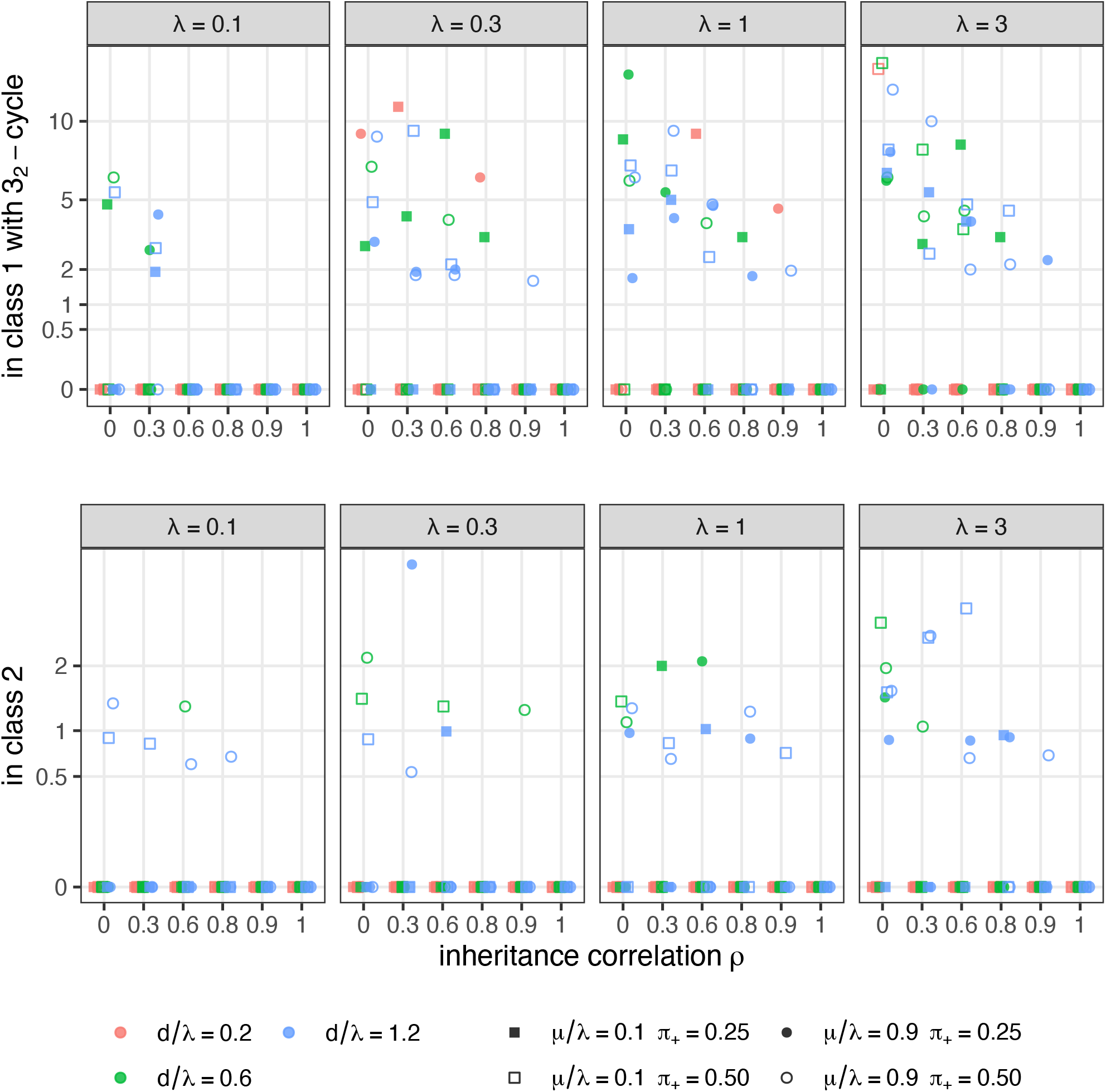
Percentage of anomalous 4-taxon networks within each subset in Fig. 6: among class-1 networks containing a 3_2_-cycle (top) and among class-2 networks (bottom). Point colors, fill and shapes are as in Fig. 8. Each point represents a variable number of four-taxon subnetworks, most influenced by *d/λ*: the sample size had an average between 11 (when *d/λ* = 0.2) and 46 (when *d/λ* = 1.2) for class-1 networks with 3_2_-cycles, and between 21 (when *d/λ* = 0.2) and 116 (when *d/λ* = 1.2) for class-2 networks.

## Appendix A. The root is not identifiable from quartet CFs

### Alternate proof of Proposition 3

Let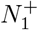 and 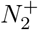 on taxa *a, b, c, d* have the same semidirected network *N* ^*−*^. Note that on 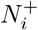 the first coalescence that occurs below the LSA determines the gene quartet tree that forms, and conditional on no coalescence occurring below the LSA each quartet forms with probability 1/3. This implies that the structure above the LSA has no effect on gene quartets, and the qCF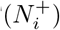 is unchanged by removing all edges and nodes above the LSA and rerooting at the LSA. We thus assume the 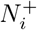 are LSA networks. However, 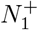 and 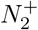 may still be rooted differently.

Let *B*_*i*_ denote the collection of edges and nodes in 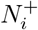 that are below or equal to at least one hybrid node of 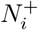. Let *H*_*i*_ be the set of hybrid nodes of 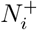, and *A*_*i*_ the remaining edges and nodes of 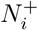. Since the semidirected networks are the same, with the obvious identifications, *B*_1_ = *B*_2_ = *B* as directed graphs, and *H*_1_ = *H*_2_ = *H*. Moreover, by making distinct copies 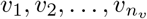 of each hybrid node *v ∈ H*_*i*_ to serve as child nodes for the *n*_*v*_ edges parental to *v*, the sets *A*_1_, *A*_2_ can be viewed as forming rooted trees, which differ only in their root locations. With this decomposition, 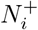 is recovered from *A*_*i*_ and *B* by identifying all the *v*_*i*_ with *v* (as is done in practice via a shared label in extended Newick notation (Cardona et al., 2008)). Let 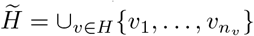.

Using that the first coalescent event to occur among four gene lineages determines the quartet tree, we get 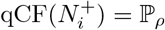 (a coalescence occurs in *B*_*i*_) qCF(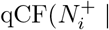| a coalescence occurs in *B*_*i*_) 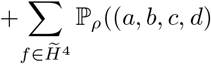 lineages reach nodes *f* with no coalescence) qCF(*A*_*i*_|(*a, b, c, d*) lineages distinct at *f*) .

Since *B*_*i*_ = *B*, the first term here is independent of *i*, as is the first factor in each term of the sum. Since *ρ* is irrelevant on a tree, and since the location of the root in tree *A*_*i*_ does not matter for quartet CFs (Allman et al., 2011), the second factor is also independent of *i*. Thus the 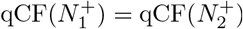.

## Appendix B. Lack of anomalies from degree-2 blobs

We prove here a weak version of Conjecture 6, under additional assumptions.

### Lemma 13.

*Let N* ^+^ *be a 4-taxon rooted metric network with a single non-trivial blob G, which is a 2-blob trapping the root with boundary {u, v}. Let a*_1_, *a*_2_ *(resp. b*_1_, *b*_2_*) be the taxa descended from u (resp. v) as in Conjecture 6*.

*Let F*_*u*_ *(resp. F*_*v*_*) be the funnel defined as the set of edges and nodes ancestral to u but not v (resp. v but not u) in N* ^+^. *Define G*_0_ *as the core subgraph of edges and nodes ancestral to both u and v, such that G is partitioned as G*_0_ ⊔ *F*_*u*_ *⊔ F*_*v*_. *Attachment nodes c*_1_, …, *c*_*m*_ *are defined as nodes in G*_0_ *that are incident to some edge in F*_*u*_ *or F*_*v*_. *If there are m ≤* 2 *attachment points and if G*_0_ *does not trap the root, then* 𝕡_*ρ*_(*a*_1_*a*_2_|*b*_1_*b*_2_ |*N* ^+^) *≥* 1*/*3, *so N* ^+^ *is* not *anomalous*.

Note that this lemma could be used recursively, to extend the result to a larger class of networks. If *G*_0_ traps the root, we can form a new network *Ñ*^+^ with fewer edges than *N* ^+^ by considering *G*_0_ as a degree-2 blob with boundary nodes *ũ* = *c*_1_ and 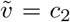 to which 2 pairs of leaves are attached. If *Ñ*^+^ is not anomalous, the proof below implies that *N* ^+^ is not anomalous either. So we can check if the funnels of *ũ* and 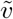 in *G*_0_ define at most 2 attachment nodes, so as to apply the lemma recursively.

*Proof*. Let *E* denote the event that a coalescent event occurs in *F*_*u*_ or *F*_*v*_, thus determining the quartet *a*_1_*a*_2_|*b*_1_*b*_2_, and 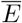 the complementary event. Let *q* be the CF of *a*_1_*a*_2_|*b*_1_*b*_2_ under NMSC(*ρ*) on *N* ^+^.

If *m* = 1 then conditioned on *E, q* = 1, while conditioned on 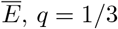, *q* = 1*/*3 by exchangeability of lineages at *c*_1_. Thus *q ≥* 1*/*3.

For *m ≥*2 note that any path from the root to *u* (resp. *v*) consists of a path from the root to some *c*_*i*_ in *G*_0_, then a path from *c*_*i*_ to *u* (resp. *v*) in *F*_*u*_ (resp. *F*_*v*_). Denote by *γ*_*i*_ (resp. *δ*_*i*_) the probability under 𝕡_*ρ*_ of the path from *c*_*i*_ to *u* in *F*_*u*_ (resp. to *v* in *F*_*v*_), which may be 0 if *c*_*i*_ is incident to *F*_*v*_ but not *F*_*u*_. Let ***γ*** = (*γ*_1_, *· · ·, γ*_*m*_) and ***δ*** = (*δ*_1_, *· · ·, δ*_*m*_). Conditioned on E, *q* = 1. Therefore

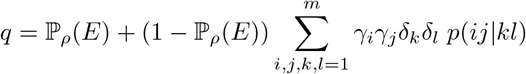

where *p*(*ij*|*kl*) is defined as a concordance factor on *G*_0_ with the attachment nodes considered as leaves:

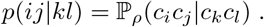

These quartet probabilities could be complicated but they only depend on the core *G*_0_, not on the funnels or the attachment probabilities ***γ*** and ***δ***. In what follows we assume that all edges have length 0 in and below *F*_*u*_ and in and below *F*_*v*_, so that coalescences do not occur between *a*_1_, *a*_2_ (or between *b*_1_, *b*_2_) before they reach the core. Indeed, this scenario minimizes *q* if all other parameters are held fixed, because it implies that 𝕡_*ρ*_(*E*) = 0 .

Now suppose that *m* = 2. Define *n*(*i, j, k, l*) ∈{2, 3, 4} as the maximum number of indices in (*i, j, k, l*) that are equal to each other. For example, *n*(1, 1, 1, 1) = 4, *n*(1, 1, 1, 2) = 3 and *n*(1, 1, 2, 2) = *n*(1, 2, 1, 2) = 2. By exchangeability, *p*(*ij*|*kl*) = 1*/*3 if *n*(*i, j, k, l*) is 3 or 4. Let *q*_0_ = *p*(11|22) = *p*(22|11). Since *G*_0_ does not trap the root, *q*_0_ *≥* 1*/*3 by Theorem 4. Then *p*(12|12) = *p*(12|21) = *p*(21|12) = *p*(21|21) = (1 *− q*_0_)*/*2 because for any *i, j, k, l* we have that *p*(*ij*|*kl*) + *p*(*ik*|*jl*) + *p*(*il*|*jk*) = 1. So we can write

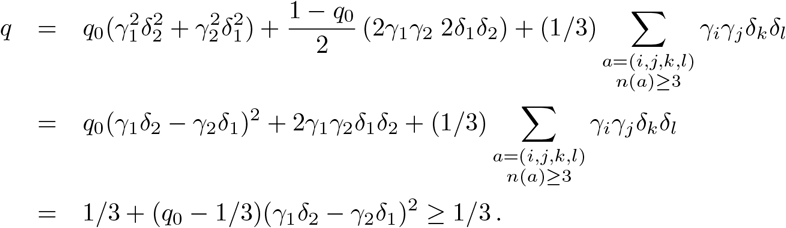

